# A complex *Plasmodium falciparum* cryptotype circulating at low frequency across the African continent

**DOI:** 10.1101/2024.01.20.576496

**Authors:** Olivo Miotto, Alfred Amambua-Ngwa, Lucas Amenga-Etego, Muzamil M Abdel Hamid, Ishag Adam, Enoch Aninagyei, Tobias Apinjoh, Gordon A Awandare, Philip Bejon, Gwladys I Bertin, Marielle Bouyou-Akotet, Antoine Claessens, David J Conway, Umberto D’Alessandro, Mahamadou Diakite, Abdoulaye Djimdé, Arjen M Dondorp, Patrick Duffy, Rick M Fairhurst, Caterina I Fanello, Anita Ghansah, Deus Ishengoma, Mara Lawniczak, Oumou Maïga-Ascofaré, Sarah Auburn, Anna Rosanas-Urgell, Varanya Wasakul, Nina FD White, Jacob Almagro-Garcia, Richard D Pearson, Sonia Goncalves, Cristina Ariani, Zbynek Bozdech, William Hamilton, Victoria Simpson, Dominic P Kwiatkowski

## Abstract

The population structure of the malaria parasite *Plasmodium falciparum* can reveal underlying demographic and adaptive evolutionary processes. Here, we analyse population structure in 4,376 *P. falciparum* genomes from 21 countries across Africa. We identified a strongly differentiated cluster of parasites, comprising ∼1.2% of samples analysed, geographically distributed over 13 countries across the continent. Members of this cluster, named AF1, carry a genetic background consisting of a large number of highly differentiated variants, rarely observed outside this cluster, at a multitude of genomic loci distributed across most chromosomes. At these loci, the AF1 haplotypes appear to have common ancestry, irrespective of the sampling location; outside the shared loci, however, AF1 members are genetically similar to their sympatric parasites. AF1 parasites sharing up to 23 genomic co-inherited regions were found in all major regions of Africa, at locations over 7,000 km apart. We coined the term *cryptotype* to describe a complex common background which is geographically widespread, but concealed by genomic regions of local origin. Most AF1 differentiated variants are functionally related, comprising structural variations and single nucleotide polymorphisms in components of the MSP1 complex and several other genes involved in interactions with red blood cells, including invasion and erythrocyte antigen export. We propose that AF1 parasites have adapted to some as yet unidentified evolutionary niche, by acquiring a complex compendium of interacting variants that rarely circulate separately in Africa. As the cryptotype spread across the continent, it appears to have been maintained mostly intact in spite of recombination events, suggesting a selective advantage. It is possible that other cryptotypes circulate in Africa, and new analysis methods may be needed to identify them.

## INTRODUCTION

The protozoan parasite *Plasmodium falciparum*, prevalent in tropical regions and especially in Sub-Saharan Africa (SSA), is responsible for causing the most severe forms of malaria, resulting in hundreds of thousands of deaths yearly.^1,2^ *P. falciparum* has repeatedly shown a great ability to adapt through genetic changes in response to human interventions, often undermining malaria control and elimination efforts.^3^ The availability of high-throughput genome sequencing in recent years has made it possible to study such changes in near-real time, providing important insights into the dynamics of genetic evolution at the population level.^4-7^ In particular, investigations into *P. falciparum* population structure– the differences in the distribution of genetic variation between populations– have revealed valuable insights into *P. falciparum* demography, by identifying patterns associated with deviations from random mating.

In highly endemic areas where malaria transmission is high, the large parasite population and frequent infection rate provide frequent opportunities for genetically distinct parasites to mate in the mosquito vectors, maintaining high levels of genetic variation through outbreeding. Hence, genetic distances within high-transmission populations tend to be evenly distributed, and no significant population structure emerges, as seen in parts of Africa.^8^ In areas where malaria transmission is low, on the other hand, parasite-carrying mosquitoes are frequently infected from biting a single individual, which results in mating between clones with identical genomes, or *selfing*. High levels of selfing result in inbred populations, which can be detected from population genomics analyses, since they exhibit lower genetic distances between individuals. High levels of inbreeding may also occur when selfing is advantageous for parasite survival. For example, a parasite that has acquired a single advantageous mutation, such as a variant that confers antimalarial drug resistance, will on average propagate this variant to only half of its offspring when mating with a wild-type. Selfing, on the other hand, guarantees that all the offspring are endowed with the advantageous variant, increasing their chances of survival, to the benefit of the whole population. Population structure driven by drug-resistant mutations was observed in the Greater Mekong Subregion (GMS), when highly inbred artemisinin-resistant populations were identified in Cambodia, each associated to a mutation in the *kelch13* gene.^9,10^

The benefits of high selfing rates are even greater when transmitting complex genetic backgrounds, as might be the case of a drug-resistant mutation that is detrimental to parasite development unless accompanied by multiple compensatory mutations. In the case of artemisinin resistance in the GMS, at least five loci were found to be co-inherited with the key *kelch13* mutations.^10^ Due to recombination, a greater number of co-inherited loci determines a lower probability of passing a complete set of variants to offspring when recombining with wild-type parasites, and therefore greater disadvantages from random mating. If offspring that inherit the full set of genetic variants are more likely to survive, then drug pressure may select lineages from selfing parasites, resulting in a reduction of genetic variation detectable by population structure analysis.

Analyses of population structure in sub-Saharan Africa have shown that high levels of genetic variations are present across high-transmission regions, with gradual genetic differentiation as one moves between East and West Africa.^11^ At the margins of malaria endemicity, however, in region with lower transmission levels such as the Gambia and the Horn of Africa, marked population structure has been observed, reflecting geographical isolation or selective pressures.^11,12^ To date, however, no published analyses have reported population structure driven by the selection of complex co-inherited multi-locus genetic backgrounds.

Here, we conducted an analysis of 4,376 African genomes from the MalariaGEN Pf7 dataset^13^ to search for patterns of population structure. By applying methods based on both *identity by state* (IBS) and *identity by descent* (IBD), we characterized a group of parasites, labelled AF1, which share a complex multi-locus genetic background, whose components appear to be co-inherited. AF1 parasites were found at very low frequency in many countries in SSA, ranging from Mauritania to Madagascar. Outside the co-inherited loci, these parasites exhibit geographical differentiation, suggesting that AF1 frequently outbreeds with local populations. We defined the term *cryptotype* to describe the characteristics of AF1, reflecting the fact that the relationship between its members is not manifest unless specific analytical methods are applied. Interestingly, the genetic background consists of common polymorphisms in genes related to parasite pathophysiology, including host-parasite interactions (antigenic variation and invasion), transmission (gametocytogenesis and mosquito stage development) and intraerythrocytic growth (gene expression regulation and biosynthesis). This suggests that this newly discovered *P. falciparum* AF1 cryptotype reflects a unique physiological or epidemiological condition coexisting within current malaria transmission through the African continent. These results provide a new perspective on evolution of malaria parasites and their transmission.

## RESULTS

### Population structure analysis of African *P. falciparum* parasites

We carried out population structure analyses on a set of African genomes from the MalariaGEN *Plasmodium falciparum* Community Project V7.0 dataset.^13^ We selected samples that were: collected in an African country; included in the quality-filtered V7.0 analysis set; and essentially clonal (i.e. had *F*_*WS*_ ≥ 0.95). This produced an analysis dataset of 4,376 from 21 countries, which were organized into three macro-regions: West, Central and East Africa-labelled WAF, CAF and EAF respectively (Table 1). Using this sample set, we estimated allele frequencies in WAF, CAF and EAF for all the high-quality biallelic single nucleotide polymorphisms (SNPs) identified in the V7.0 dataset. We removed extremely rare variants by discarding those SNPs whose minor allele frequency (MAF) was <0.1% in all three major populations, resulting in a set of 743,584 high-quality SNPs.

**Table 1.** Summary of sample counts by country. Each row represents one African country where *P. falciparum* samples analysed in this study were samples. The columns show: the macro-region in which the country is located (West, Central or East Africa); the name of the country and its ISO 3166 code; the total number of analysed samples from that country; the number of AF1 samples identified in the country, and their percentage of the samples analysed.

| Macro-region | Country Name | Country Code | Sample Count | AF1 Count | AF1 % |
| --- | --- | --- | --- | --- | --- |
| West Africa (WAF) | Mauritania | MR | 53 | 1 | 1.9% |
|  | Mali | ML | 627 | 4 | 0.6% |
|  | Senegal | SN | 112 |  |  |
|  | Gambia | GM | 530 | 3 | 0.6% |
|  | Guinea | GN | 76 | 5 | 6.6% |
|  | Ghana | GH | 1,430 | 19 | 1.3% |
|  | Ivory Coast | CI | 45 | 1 | 2.2% |
|  | Burkina Faso | BF | 17 |  |  |
|  | Benin | BJ | 93 |  |  |
|  | Nigeria | NG | 78 |  |  |
|  | Cameroon | CM | 128 |  |  |
| Central Africa (CAF) | Gabon | GA | 33 | 1 | 3.0% |
|  | DR Congo | CD | 233 | 1 | 0.4% |
| East Africa (EAF) | Sudan | SD | 64 |  |  |
|  | Ethiopia | ET | 19 |  |  |
|  | Kenya | KE | 379 | 1 | 0.3% |
|  | Uganda | UG | 5 | 1 | 20.0% |
|  | Tanzania | TZ | 315 | 4 | 1.3% |
|  | Malawi | MW | 102 | 5 | 4.9% |
|  | Mozambique | MZ | 17 |  |  |
|  | Madagascar | MG | 20 | 1 | 5.0% |
| Total |  |  | 4,376 | 47 | 1.2% |

We computed a matrix of pairwise genetic distances between all samples, to which we applied principal coordinate analysis (PCoA), a method that maps samples onto a series of dimensions (*principal components*) that explain matrix variance; when plotting principal components, clusters of genomes that are highly similar to each other (e.g. inbred populations) tend to separate from other individuals. A plot of the first two components, which explain the most variance, is shown in Figure 1. The first component (PC1) was driven by the differentiation between WAF and EAF parasites, with CAF populations occupying a middle space; such differentiation has been shown in previous reports.^4,11^,14 Higher-order components (PC3 to PC7), on the other hand, appeared to be driven by clusters of parasites from Ethiopia and The Gambia (Supplementary Figure 1), consistent with population structure in low-endemic regions, described in previous work.^11^ The second component (PC2) was also driven by a diverging cluster, which we named AF1, which was composed of parasites sampled from multiple countries across Africa, rather than from sites in close geographic proximity.

**Figure 1.**
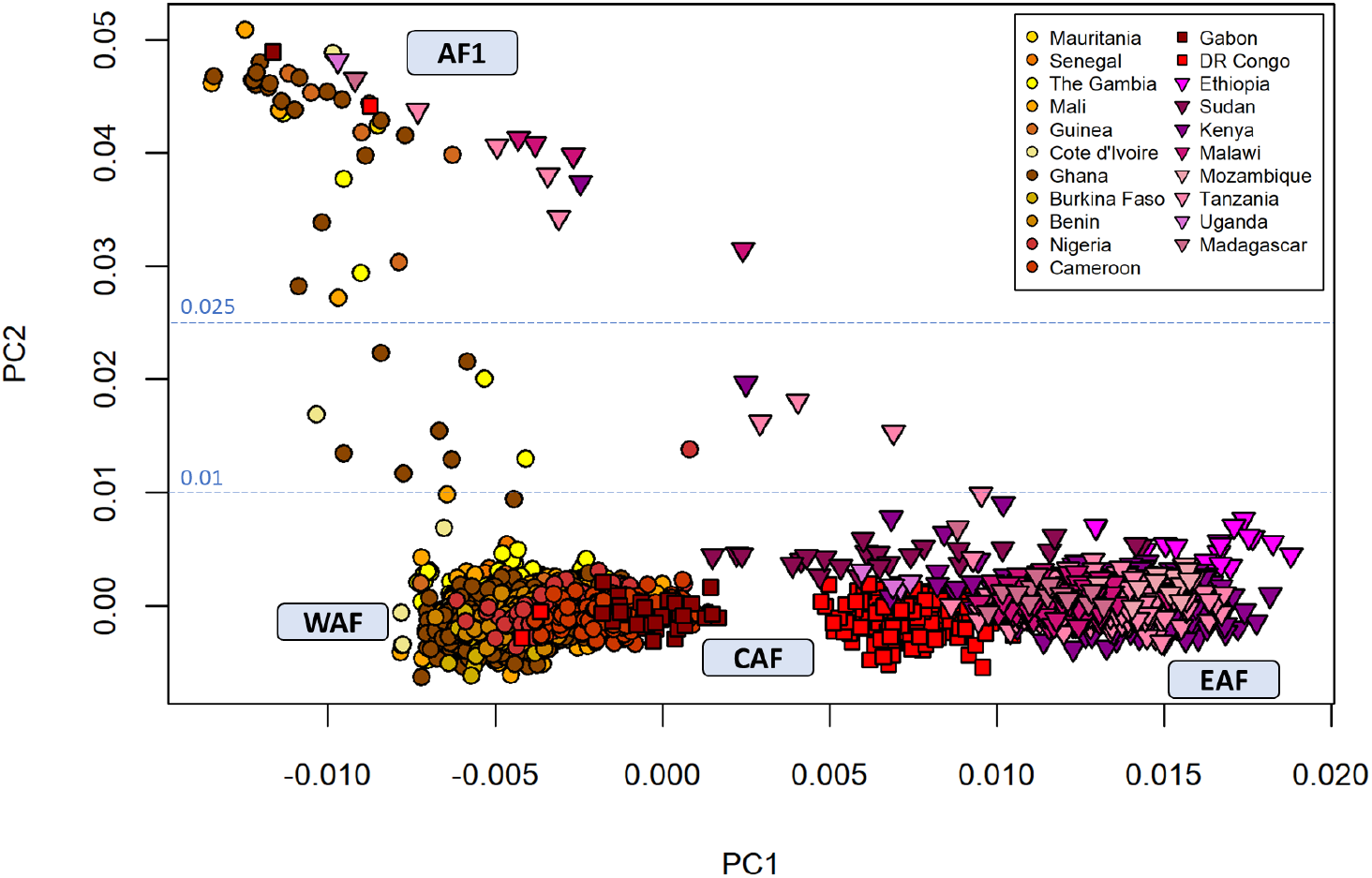
Principal Coordinate Analysis (PCoA) of African samples, revealing population structure. In this figure, we show plots of the second principal component against the first (PC2 vs PC1). Along PC1, samples separate geographically, so that macro-regions EAF, CAF and WAF can be distinguished as labelled. A cluster of AF1 parasites, from different countries, separates along PC2. Two horizontal dotted lines indicate the thresholds for defining the AF1 population. Samples with PC2 > 0.025 were classified as AF1; those with PC2 < 0.01 as non-AF1; the remaining parasites were disregarded in further analysis, since their AF1 membership status is uncertain.

The broad geographical distribution of AF1, which includes regions of high transmission, sets this cluster apart from those identified by higher-order components, and rules out population structure driven by low endemicity. Rather, it suggests that AF1 members share a high degree of similarity in a substantial portion of the genome, and this differentiates them from other individuals collected from the same countries.

We labelled samples with PC2 ≥ 0.025 as AF1 members (Figure 1), while parasites with PC2 ≤ 0.01 were assigned labels according to their macro-region (WAF, CAF or EAF); samples with intermediate PC2 values (n=14) were disregarded. The AF1 group comprised 47 samples from 13 countries in all three macro-regions, sampled up to 7,500 km apart from each other (Figure 2). Overall, AF1 parasites represent about 1.2% of all samples in the analysed dataset (Table 1). Within most countries, AF1 accounts for 1-6% of samples, and there is no significant difference between the AF1 frequencies in the two most represented macro-regions (1.2% in WAF, 1.46% in EAF, *p*=0.547). Thus, to a first level of approximation, AF1 appears to be fairly evenly distributed, at low frequency across the continent.

**Figure 2.**
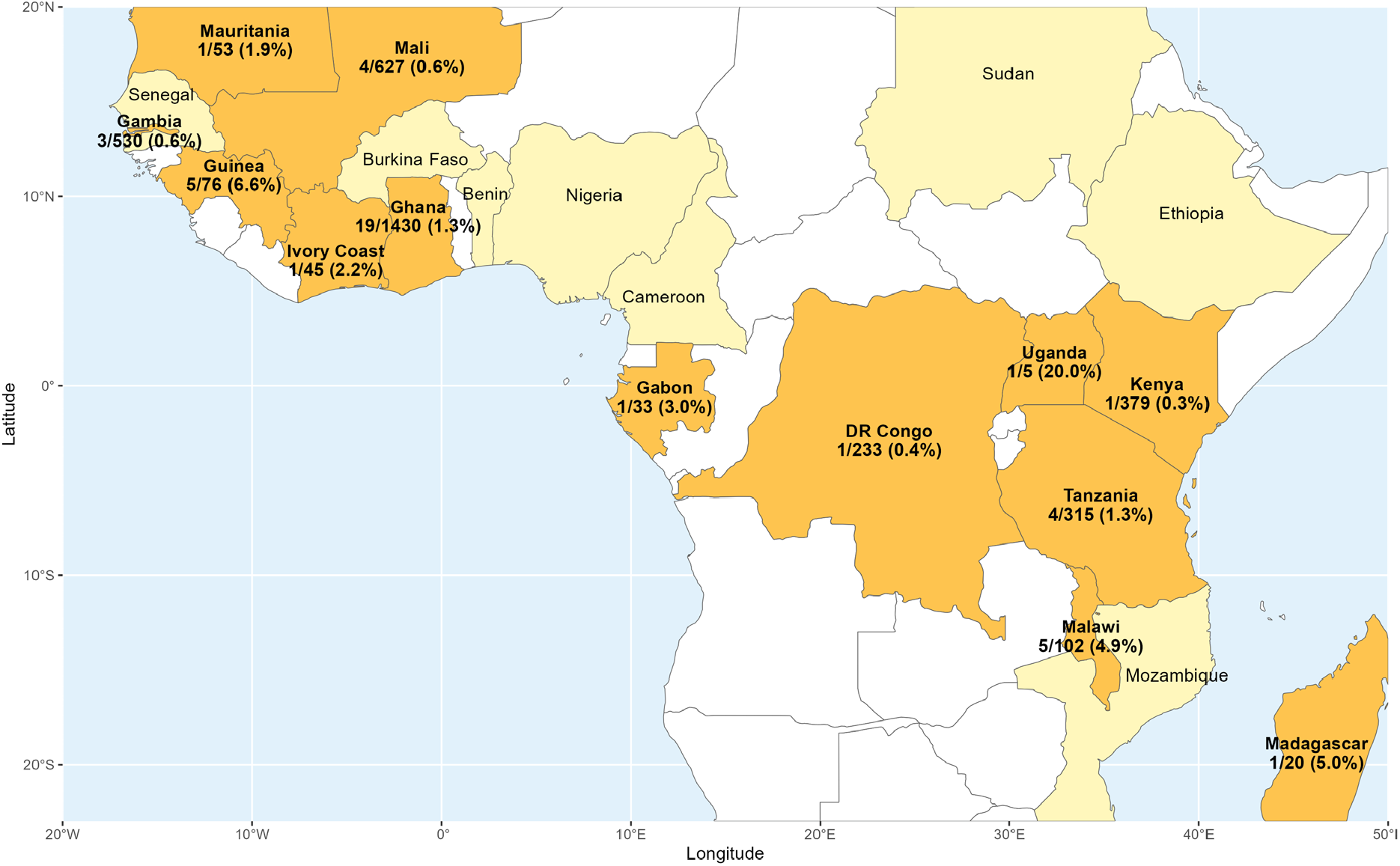
Geographic distribution of AF1 parasite samples. In the map above, countries from which parasites were sampled are shown with a coloured background and a label showing the country name. Countries where AF1 parasites were found are shown with an orange background; for each of these countries, the number of AF1 samples and the total number of analyzed samples are separated by a slash, and the percentage of AF1 samples is shown in brackets. The map uses data from Natural Earth (https://www.naturalearthdata.com/)

### Genetic features of AF1

The clustering of AF1 parasites suggests that they share alleles in common that are not common in the rest of the African populations. To identify genetic loci that differentiate AF1, we estimated allele frequencies in AF1, WAF, CAF and EAF at all coding SNPs, and used these frequencies to estimate the mean *F*_*ST*_ between AF1 and each of the other three populations. For this task, we were able to include 68,360 additional SNPs that exhibited high missingness in samples processed using selective whole-genome amplification (sWGA) but were well-covered in non-amplified samples (n=1,829 in WAF, CAF and EAF).

This analysis revealed a large number of highly differentiated loci in gene exons: we found 199 coding non-synonymous SNPs with mean *F*_*ST*_ ≥0.5, and 69 of these had mean *F*_*ST*_ ≥0.75 (Supplementary Table 1). When mapping the most differentiated SNPs across the genome, they do not appear evenly distributed, but rather clustered in several regions of the genome, on multiple chromosomes (Supplementary Figure 2). In particular, we found cluster of high-*F*_*ST*_ variants in chromosomes 1, 2, 4, 9, 10, 11, 13 and 14, while other chromosomes showed lower levels of differentiation. The presence of clusters of high-*F*_*ST*_ SNPs in close proximity suggests that some of these *AF1 characteristic loci* contain highly differentiated long haplotypes. Although most of the SNP clusters occupy regions <100kbp, one large set of highly differentiated SNPs on chromosome 10 stretches over ∼250kbp, possibly indicating a haplotype under selection, or a large structural variant.

Given the high degree of differentiation at the AF1 characteristic loci, we predicted a high degree of correlation between SNPs located in these regions. This was confirmed by computing *r*^*2*^, a commonly used measure of linkage disequilibrium,^15^ for all distal pairs of SNPs with moderate-to-high AF1 differentiation (mean *F*_*ST*_≥0.2). Several loci were found to contain SNP highly correlated with distal SNPs (*r*^*2*^≥0.2); mapping these associations across the genome produces a complex linkage network (Figure 3). Six of these loci formed a network in which at least one SNP at each locus showed very strong association (*r*^*2*^ ≥0.4) with SNPs at all other loci (Supplementary Table 2). This provides clear evidence that AF1 parasites possess a multi-component genetic background, carried as a complete set by most members. However, the exact composition of this background cannot be fully determined with this analysis, since high *r*^*2*^ values are only possible when the AF1 characteristic alleles are very rare outside AF1, which is not a requisite for a component locus.

**Figure 3.**
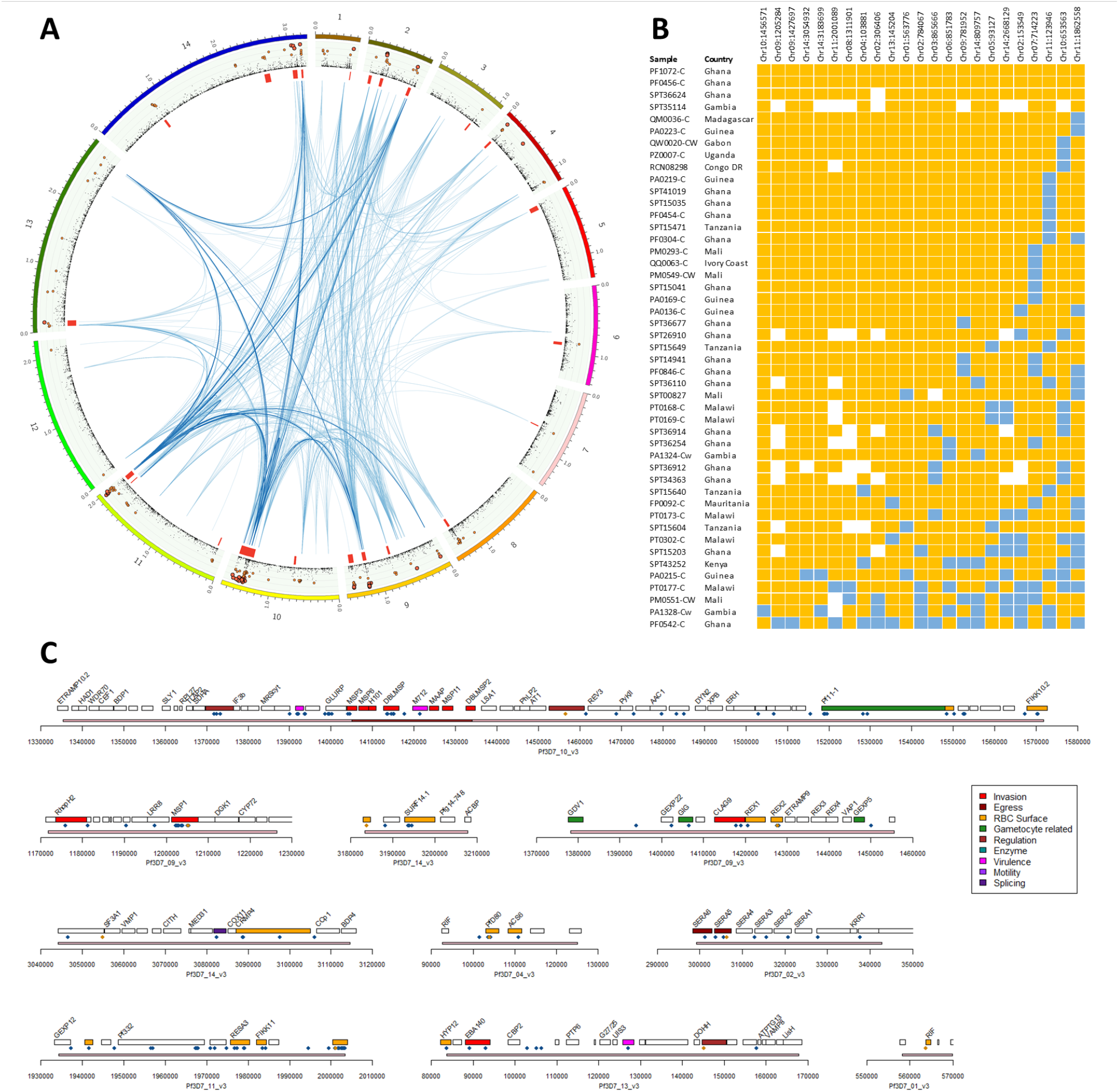
A – Linkage between AF1 characteristic loci. The circular plot maps all 14 nuclear chromosomes (starting from top, clockwise, each chromosome is represented by a coloured segment in the outer ring). The ring with a green background shows a plot of mean *F*_*ST*_ between AF1 and the three African macro-regions (WAF, CAF and EAF) at non-synonymous coding SNPs with *F*_*ST*_ ≥0.01, with orange markers showing *F*_*ST*_ ≥0.5 and larger red markers indicating *F*_*ST*_ ≥0.75 (see Supplementary Table 1). Inside this ring, red strips delimit high-IBD regions, in which ≥50% of all AF1 sample pairs are in IBD (see Supplementary Figure 5). Inner blue lines show the *r*^*2*^ measure of linkage disequilibrium (LD) between pairs of high-*F*_*ST*_ SNPs (*F*_*ST*_ >0.2), estimated using all African parasites. Three types of line represent three LD ranges: 0.2 ≤ *r*^*2*^ <0.4 (lightest blue, thinnest lines), 0.4 ≤ *r*^*2*^ <0.5, and *r*^*2*^ ≥0.5 (deepest blue, thickest line). **B – Presence of characteristic haplotypes in AF1 parasites**. This panel shows a matrix of genotypes at the 23 SNPs with the highest *F*_*ST*_ in the 23 high-IBD regions identified in the AF1 population. Each row represents an AF1 sample; the sample ID and the country of provenance are shown. Blue cells indicate that the sample carried the reference allele, while orange cells indicate the characteristic AF1 allele (non-reference). White cells indicate a missing genotype. **C – Genes at AF1 characteristic loci**. This panel shows maps of gene positions for the 10 highest-ranked high-IBD regions identified in the AF1 population. The x-axis represents positions on the high-IBD region’s chromosome, with a pink line showing the extent of the region. Each gene in the region is shown by a rectangle, labelled with the gene’s name and coloured according to its function (where they are known). Non-synonymous coding SNPs with mean *F*_*ST*_ ≥0.5 and shown by blue diamond markers below the gene boxes; the highest-*F*_*ST*_ SNP in each region (see Supplementary Table 2) is denoted by an orange marker.

### Ancestry analysis

Given the broad geographic distribution of AF1 members and the complexity of their common genetic background, we investigated whether their shared alleles originate from different sources in different countries, or they are likely to have been co-inherited from common ancestry. To address this question, we conducted an analysis of identity by descent (IBD) for all sample pairs in our dataset. This analysis identifies stretches of the genome where a pair of parasites shows a degree of similarity that is unlikely to be observed unless the two parasites have a common ancestry. The AF1 parasites exhibited pairwise IBD at a much higher fraction of their genomes (median=22.6%) than non-AF1 parasites in West, Central and East Africa (medians=0.0%, 0.8% and 1.1% respectively, Supplementary Figure 3A), suggesting that the genomic regions that differentiate AF1 have common ancestries. This is confirmed when analysing PCoA using a distance measure derived from IBD genome fractions, which shows AF1 samples forming a highly compact outlier group (Supplementary Figure 4). Even though the pairwise IBD levels are well above the average in African populations, AF1 does not have the characteristics of a clonally expanding population that relies on selfing. West African AF1 genomes shared significantly higher IBD fractions with West African non-AF1 genomes than with East African ones (median=0.56% vs 0.12%), and vice versa (median=0.55% vs 0.0%, Supplementary Figure 3B), indicating that recombination occurs between AF1 parasites and non-AF1 local populations.

Hypothesizing that IBD in AF1 samples is restricted to specific regions, we mapped the frequency of IBD segments in AF1 pairs across the genome, identifying 23 “high-IBD” regions where at least 50% of all AF1 pairs were in IBD (Supplementary Figure 5). High-IBD regions were present in all chromosomes except chromosome 12, and in several cases appeared to be located near subtelomeric regions. Remarkably, every high-IBD region contained one or more SNPs with mean *F*_*ST*_>0.5 and the AF1 allele frequency >0.5 (Supplementary Table 3). When inspecting sample genotypes at these 23 loci, it appears that the high-*F*_*ST*_ SNPs, ranked by allele frequency, are effective markers for identifying AF1 members: most samples (42/47) carry the AF1 characteristic alleles at all top 7 ranked SNPs, and no more than one non-AF1 allele in the top 13 SNPs (Figure 3B). Conversely, AF1 characteristic alleles were rare in non-AF1 genomes: only one out of 4,326 samples carried AF1 alleles at more than half of the six top ranked SNPs, suggesting that AF1 members can be discriminated by means of relatively simple genetic tests.

When putting together results from analyses of IBD, differentiation and correlation, we find substantial overlap: highly differentiated loci are mostly located in regions of high IBD and strongly linked across chromosomes (Figure 3A). Hence, we can deduce that AF1 parasites carry a constellation of numerous variants that differentiate them from the rest of African parasite, and appear to be inherited together, even though AF1 genomes evidently undergo recombination with sympatric strains. It appears that not all the loci involved are equally important; a “core” set of around 13 loci is present in most AF1 members, with very few discordances; while other loci seem to be less critical members of the constellation. All evidence suggests that the constellation of variants is co-inherited, rather than come from recombination of individuals with different ancestries in different countries.

### Structural variants in chromosome 10 and 9

The top-ranked high-IBD region, on chromosome 10 (Figure 3C), is considerably larger than all other high-IBD regions, and unlikely to be underpinned by a single SNP variant; hence, we hypothesized that it may harbour a structural variant. This region contains several related members of the MSP3 gene family, encoding merozoite surface proteins involved in erythrocyte invasion.^16^ A sequencing read coverage analysis found that AF1 members had few or no reads mapping to genes MSP6 (PF3D7_1035500) and H101 (PF3D7_1035600), suggesting a large deletion (Supplementary Figures 6 and 7). The adjacent 5’ end of the DBLMSP gene (PF3D7_1035700) was also poorly covered, but this is more difficult to interpret due to the presence of a paralog of DBLMSP (DBLMSP2, PF3D7_1036300) which is a possible cause of read mismapping. To get a clearer picture, we performed *de novo* assembly of the sequencing reads of an AF1 member from Mali (PM0293-C) and mapped the resulting contigs to multiple reference sequences of the DBLMSP and DBLMSP2 genes, provided by the Pf3k project (https://www.malariagen.net/projects/pf3k).^17^ The DBLMSP sequence of AF1 shows a high degree of similarity to the PfIT reference (a South American strain), but a very different organization (Figure 4A). While the AF1 sequence is almost identical to the PfIT DBLMSP sequence downstream of position 1010, at the 5’ end it is almost identical to the PfIT DBLMSP2 gene. This suggests that the AF1 DBLMSP gene has undergone gene conversion, acquiring a portion of a DBLMSP2 gene at the 5’ end; we could not establish whether this occurred in the same recombination event that deleted the neighbouring MSP6 and H101 genes. At the join between the two gene portions, we identified a 19-nt stretch which is identical in DBLMSP and DBLMSP2, and likely to be the recombination breakpoint (Figure 4B). A 62-nt sequence, comprising the breakpoint segment and flanking fragments, was used as a search template to confirm the presence of the recombinant DBLMSP by inspecting the sequencing reads of AF1 members. Matches for the sequence were found in 42 out of 47 AF1 samples; in the remaining samples we could not identify the DBLMSP sequence, possibly because of an alternative structural variant, or because of localized poor coverage. The DBLMSP2-like sequence at the 5’ end of the AF1 DBLMSP explains the absence of coverage when aligning AF1 reads against the 3D7 reference; a realignment using the *de novo* assembled AF1 sequences as references showed elevated coverage at the 5’ end, confirming that both AF1 genes possessed DBLMSP2-like sequences in that regions (Supplementary Figure 8). Although DBLMSP/DBLMSP2 gene conversions have been recently reported,^18^ the AF1 conversion presents a different recombination pattern. The AF1 DBLMSP2 gene did not show signs of conversion, but showed greater similarity to the PfKH02 reference strain (from Cambodia) than to PfIT. It should be noted that other genes in this region contain additional AF1-differentiated SNPs, including MSP3 (PF3D7_1035400) and the glutamate-rich protein GLURP (PF3D7_1035300).

**Figure 4.**
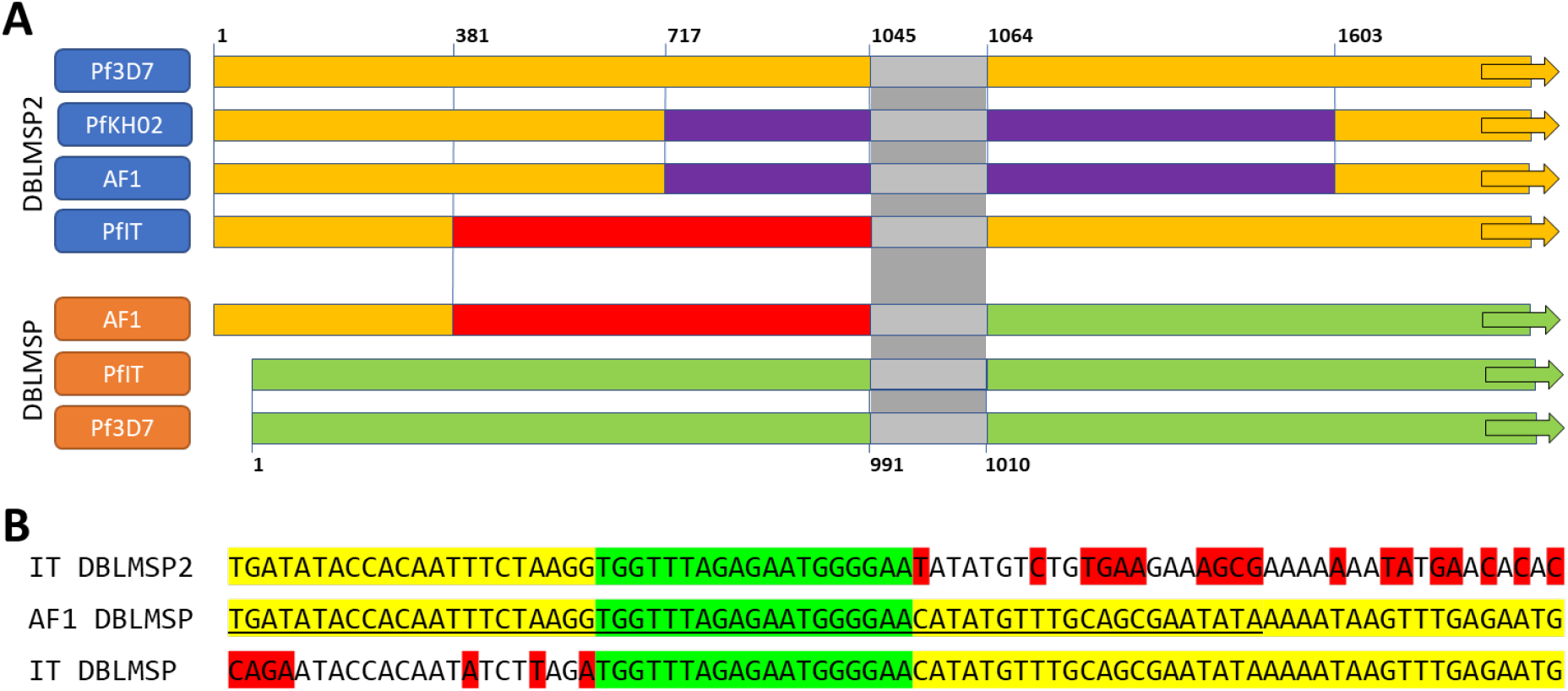
DBLMSP gene sequence crossover in AF1 parasites. **Panel A – Schematic of the gene conversion underpinning the AF1 variant of DBLMSP.** The diagram shows as colour blocks the sequences of DBLMSP and DBLMSP2 in four Pf genomes: Pf3D7 (reference), PfIT, PfKH02 (both long-read sequenced) and AF1 (*de novo* assembly of sample PM0293-C). Blocks of the same colour indicate highly similar (near-identical) sequences. Coordinates shown (not to scale) correspond to the Pf3D7 positions in DBLMSP2 (above) and DBLMSP (below). The AF1 DBLMSP sequence is near-identical to that of PfIT DBLMSP2 at the 5’ end, and of PfIT DBLMSP after position 991. The AF1 DBLMSP2 sequence, on the other hand, is near-identical to the DBLMSP2 sequence of PfKH02. The gray region is a 19-nt sequence identical in DBLMSP and DBLMSP2, likely to be the recombination breakpoint. **Panel B – Detail of the AF1 DBLMSP/DBLMSP2 breakpoint region**. This panel shows an alignment of the AF1 DBLMSP sequence (middle) against the DBLMSP2 (above) and DBLMSP (below) sequences of PfIT. The 19-nt region of 100% identity is shown in green; to the left, the AF1 sequence is identical to PfIT DBLMSP2, while to the right it is identical to PfIT DBLMSP. Allele differences with respect to the AF1 sequence are highlighted in red. The underlined 62-nt portion of the AF1 sequence was used as search query to confirm the presence of the conversion breakpoint in the AF1 parasites.

The second-ranked high-IBD region, on chromosome 9 (Figure 3C), exhibits a sequence of high-*F*_*ST*_ SNPs, suggesting a highly differentiated haplotype in the merozoite surface protein MSP1 gene (PF3D7_0930300). Furthermore, some regions of this gene appear to lack coverage in the AF1 parasites (Supplementary Figure 9), suggesting that regions of the AF1 sequence differ greatly from the reference. MSP1 is known to undergo frequent recombination at multiple sites^,19,20^ and frequently classified based on variants present in four sections of the gene.^21^ We aligned contigs from the *de novo* assembly of PM0293-C against sequences of the MAD20, K1 and RO33 strains that are used for genotyping MSP1.^22^ PM0293-C was found to carry a MAD20/K1/K1/K1 MSP1 signature; its MSP1 variant is uncommon in Africa outside the AF1 population, but at higher frequency in South America and Southeast Asia (Supplementary Table 1). The PM0293-C amino acid sequence is identical to that of PfHB3, a Mesoamerican strain, except for a 9-amino acid repeat insertion after PfHB3 position 86, and a deletion at positions 722-729.

### Functional analysis of AF1 characteristic loci

The complexity of the AF1 genetic background, the large number of loci involved, and the low frequency of the characteristic alleles outside the AF1 group suggest an extremely low probability that, when recombining with non-AF1 parasites, the offspring may inherit a full complement of AF1 alleles. Therefore, if these parasites are present at a detectable frequency, it is likely that possessing the complete allele constellation affords some fitness advantage, resulting in positive selection. This suggests a strong synergy between the alleles in the constellation, and a functional relationship between the genes that carry them. The loci on chromosomes 10 and 9 are known to be functionally related: MSP1 binds with multiple other surface proteins to forms a large complex on the merozoite surface,^23^ which acts as a platform for display of erythrocyte-binding domains of various antigens, including DBLMSP, DBLMSP2 and MSP6.^24,25^ The MSP1 complex has been shown to play a critical role in merozoite egress and invasion of erythrocytes.^26,27^ Also related to merozoite egress, the genes SERA5 (PF3D7_0207600) and SERA6 (PF3D7_0207500), known to play an essential role in disassembling the erythrocyte cytoskeleton and rupturing its membrane^,28,29^ carry multiple high-*F*_*ST*_ SNPs on chromosome 2. Further high-*F*_*ST*_ variants in genes involved in erythrocyte membrane interaction were found at other AF1 characteristic loci; among these are other merozoite surface proteins (MSP7 and MSP10), and at least 5 members of the PHIST gene family,^30^ including those encoding the essential proteins RESA3 (PF3D7_1149200) and PfD80 (PF3D7_0401800), the latter involved in trafficking proteins from the Maurer’s clefts to the erythrocyte membrane.^31^ Other high-*F*_*ST*_ SNPs are in genes associated with Maurer’s clefts and trafficking of exported proteins, such as the membrane associated histidine-rich protein MAHRP1 (PF3D7_1370300),^32^ the Pf332 antigen (PF3D7_1149000);^33^ and the ring-exported proteins REX1 and REX2 (PF3D7_0935900 and PF3D7_0936000 respectively).^34,35^ In addition, several genes encoding erythrocyte-exported proteins carry AF1 differentiated alleles, such as members of the FIKK family of kinases FIKK10.2 (PF3D7_1039000) and FIKK11 (PF3D7_1149300);^36^ SURFIN surface antigens SURFIN 4.2 (PF3D7_0424400), SURFIN 8.2 (PF3D7_0830800) and SURFIN 14.1 (PF3D7_1477600); and the rhoptry-localized cytoadherence-linked asexual gene CLAG9 (PF3D7_0935800).^37^ Overall, we find that high-*F*_*ST*_ variants in the AF1 characteristic regions are mostly in genes that fall into a small number of functional categories (Figure 3C), primarily related to erythrocyte invasion (e.g. MSP, SERA genes, EBA-175), to erythrocyte surface antigens (REX, FIKK, SURFIN, etc.), and to regulatory functions (e.g. CLAG, AP2-EXP, HSP70). Although at this point we cannot ascertain which of these variants SNPs are functionally important, the evidence points to a constellation of variants that are functionally linked, and related to host-parasite interactions.

## DISCUSSION

Labelling a pathogen by its “strain” is valuable device for describing the ancestry of viruses and bacteria, but is not easily applied to recombinant organisms, in which each genomic locus has a separate ancestry.^38^ *P. falciparum* “strains” have occasionally been labelled according to shared inherited genetic traits, such as drug-resistant haplotypes or specific recombination patterns, e.g. in the MSP1 gene.^22^ Usually, however, only one or two genomic loci are used for such classification. In the analyses presented here, based on more than 4,300 high-quality *P. falciparum* genomes, we have identified a genetic background of remarkable complexity, which circulates across the breadth of the African continent, and maintains its integrity without solely relying on inbreeding. To our knowledge, this is the first report of what we describe as a *cryptotype*, a complex inherited genetic background “hidden” within genomes that are otherwise similar to their sympatric parasites. Their shared ancestry patterns do not resemble those of clonally expanding populations,^7^ in that IBD is not evenly distributed across the genome, but rather concentrated in numerous physically distal regions. Co-occurring IBD regions around drug resistance-implicated loci have been previously observed,^11^ but in much smaller numbers and with weaker linkage. This cryptotype’s ability to retain a high degree of identity at its characteristic loci, in spite of recombination events over the long period of time it must have taken to achieve its geographic spread, is hard to reconcile with the extremely low probability that the entire constellation of variants would be passed down to progeny other than by selfing. It is possible that the AF1 genome has undergone structural changes that increase linkage between some of the characteristic regions, but this is unlikely to explain co-inheritance for such a high number of loci. It seems more likely that the frequency at which AF1 is found in our sample set (∼1%) is maintained by some selective advantage. In other words, the results strongly suggest that the variants in the AF1 constellation work in concert, interacting and possibly compensating for each other to confer added fitness, leading to selection of this genotype. Although not all of the ensemble variants are found in all AF1 parasites, suggesting they are not all equally essential, the fact that more than 20 identical AF1 variants are found in parasite from Madagascar, Ghana, Uganda and DR Congo indicates a complex and functionally important interplay between these loci. In spite of recombination, the AF1 characteristic variants are rarely found in the non-AF1 population, which suggests that on their own they are less beneficial, or even detrimental to parasite fitness.

Although at this point we cannot determine the functional changes conferred by the AF1 ensemble, an examination of the genes containing the most highly differentiated variants reveals major functional relationships, bolstering the hypothesis of strong interactions. Several of the genes involved encode proteins known to participate in erythrocyte egress and invasion. Notably, these include components of the MSP1 complex, a key player in the red blood cell invasion process; and the DBLMSP/DBLMSP2 proteins, which bind the MSP1 complex to the erythrocyte during invasion. In addition, several genes harbouring AF1 variants related to the export and trafficking of antigens expressed on the red blood cell surface. Taken together, these lines of evidence suggest that the advantage provided by the AF1 variant ensemble is related to interactions with host erythrocytes, and particularly with erythrocyte membranes. We hypothesize that AF1 parasites have adapted to a specific host erythrocyte phenotype, providing them with a niche within which they are particularly fit, e.g. the AF constellation may confer an adaptation to hemoglobinopathies that reduce invasion^39^ or prevent erythrocyte remodelling.^40^ While the broad geographic distribution makes it unlikely that cryptotype is fine-tuned to target a specific human population, we cannot exclude the possibility that its evolutionary niche involves a non-human host.

The present analysis opens several questions that will require further investigation in multiple directions. High-quality long-read sequencing of AF1 parasites will be needed to confirm the large variants identified in this study, and explore structural rearrangements in the AF1 genome. Culturing *in vitro* field isolates could provide an understanding of the biological mechanisms underpinning the cryptotype and the properties that confer its selective advantage. Identifying patients infected with AF1 parasites may help characterize the cryptotype’s evolutionary niche and understand its epidemiology. Given AF1’s low prevalence, such studies will be challenging, but they may produce important shifts in our understanding of invasion mechanisms, and of how protective human blood phenotypes work. The wide-ranging catalogue of potentially interacting variants we identified can already provide experimental parasitologists with a list of candidates for studying gene interactions and synergies.

A legitimate question that emerges from this work is whether AF1 is the only cryptotype present in Africa or, indeed, globally. In our analysis, AF1 parasites separated clearly in PCoA plots, partly because they share large haplotypes over occupying a significant proportion of their genome, and also because these differentiated variants are mostly absent from the rest of African parasites. This produces high levels of differentiation, which caused AF1 members to separate from other African parasites. However, neither the number of loci nor their size, nor their absence from the general population, are requisites for a cryptotype. Clusters of individuals could be circulating, carrying co-inherited variant sets that are harder to detect, for example because the variants are not uncommon outside the clusters. To identify such clusters, PCoA may be a blunt tool, and new approaches may be needed, based on more sensitive IBD detection algorithm and higher-resolution genotypes.

Furthermore, it is possible that other cryptotypes circulate at a low frequency like AF1, or lower, and their detection will require increasingly large genomic datasets. We have shown that shared genomic data, aggregated from a multitude of studies in different countries, can lead to important discoveries. The shared genomic repositories that provide such data in an organized and usable form, such as the MalariaGEN *P. falciparum* Community Project,^13^ require much effort and resources to maintain and grow. We advocate that these repositories should continue to be supported by funders and contributing researchers alike, to power our advancements towards a better understanding of epidemiological phenomena.

## METHODS

To construct an analysis dataset to investigate population structure in Africa, we began with the Plasmodium falciparum Community Project v7 (Pf7) dataset,^13^ which comprises a total of 20,864 samples. The release includes metadata and preliminary analysis results, including the identification of an “analysis set” comprising 16,203 samples with low genotyping missingness. From this set, we selected “essentially clonal” samples (*F*_*WS*_ ≥0.95, using the Pf7 release *F*_*WS*_ estimates) from Africa, resulting in a set of 4,376 samples, which were organized by macro-regions: West, Central and East Africa-labelled WAF, CAF and EAF respectively (Table 1). After estimating allele frequencies at 2,025,136 biallelic PASS SNPs where ≥75% of samples could be genotyped, we selected those SNPs whose minor allele frequency (MAF) was ≥0.1% in at least one macro-region, producing a set of 743,584 high-quality SNPs. Based on the assumption of near-clonality, all samples were genotyped at each SNP with the allele supported by the highest number of reads; hence, all genotypes were called as homozygous. Allele frequencies in each macro-region were estimated at each SNP by calculating the proportion of samples carrying each allele, disregarding samples with a missing genotype. Pairwise *F*_*ST*_ at any given SNP was estimated from the non-reference allele frequencies (NRAF) in the two populations, as previously described.^10^ The AF1 mean *F*_*ST*_ mean was calculated as the arithmetic mean of *F*_*ST*_ between AF1 and each of the macro-regions WAF, CAF and EAF.

We observed that several genome segments were poorly covered in samples processed by selective whole-genome amplification (sWGA, a laboratory protocol that enables whole-genome sequencing from dried blood spots), causing high missingness in variants within these segments when analysing the entire sample set. To obviate this problem and discover a more complete set of differentiated SNPs, we repeated the SNP filtering and *F*_*ST*_ estimation procedures on the subset of samples that were not amplified prior to sequencing (n=1,829 in WAF, CAF and EAF). This produced an additional set of 68,360 SNPs; these were only used in the identification of AF1 highly differentiated variants.

To analyse population structure, we constructed an *NxN* pairwise genetic distance matrix (*N*=4,376). Genome-wide genetic distances, based on genotypes at the 743,584 SNPs, were estimated by the proportion of SNPs where the two samples carry different alleles, after discarding SNPs where one or both of the two samples have a missing genotype. Analyses of population structure were performed using a combination of freely available tools, and bespoke software programs written in Java and R. PCoA analyses were carried out using the stats package in the R language (https://www.r-project.org/), while neighbour-joining trees were produced by the R ape package.^41^

The linkage disequilibrium measure *r*^*2*^ between two SNPs *S*_*1*_ and *S*_*2*_ was calculated from the allele frequencies of the major and minor allele at *S*_*1*_ (*p*_*1*_ and *p*_*2*_ respectively) and *S*_*2*_ (*q*_*1*_ and *q*_*2*_), and the *allele pair frequencies x*_*11*_, *x*_*21*_, *x*_*12*_, *x*_*22*_ (the indices indicating whether the major or minor allele is present at each of the two SNPs) as follows: *r*^2^ = *D*^2^/(*p*_1_*p*_2_*q*_1_*q*_2_) where *D* = *x*_11_ − *p*_1_*q*_1_.^15^ Only SNPs with mean *F*_*ST*_ ≥0.2 were used in linkage pairs. The circular genome linkage plot was generated using circos v0.69.^42^

Identity by descent (IBD) analysis was performed on the genotypes obtained from whole-genome sequencing using the program hmmIBD^43^ using default parameters. For this analysis, we filtered out variants at extremely low frequency, using coding SNPs with MAF ≥0.1 in at least one of the macro-regions, and at least one sample with a homozygous non-reference genotype. High-IBD regions in the AF1 population were defined by identifying uninterrupted sequences of SNPs where ≥50% of all AF1 pairs were in IBD. Neighbouring regions separated by gaps ≤50kbp were subsequently merged. To perform an IBD-based population structure (PCoA) analysis, we constructed an *NxN* pairwise genetic distance matrix (*N*=4,376), where distance was calculated as *d*=(1-*f*_*IBD*_) where *f*_*IBD*_ is the fraction of the genome that is predicted to be IBD in a given pair.

De novo assemblies of genomic sequencing reads were executed using Cortex v1.0.5.21^44^ with kmer_size=61. The generated contigs were aligned against reference sequences provided by the Pf3k project (https://www.malariagen.net/projects/pf3k)^17^ using BioEdit v7.2.5 (https://thalljiscience.github.io/). MSP1 gene references were obtained from GenBank, accession numbers X03371.1 (K1), AB276005.1 (RO33) and X05624.2 (MAD30).

Coverage analyses were conducted using normalized coverage at 743,584 SNPs. For each sample, we obtained the number of reads covering each SNP, determined the median coverage across the sample, and normalized coverage by dividing SNP coverages by the sample’s median coverage.

Sequencing reads coverage visualizations were produced using the LookSeq web application.^45^ Functional information about genes was obtained from the PlasmoDB database (https://plasmodb.org/plasmo/app/) and from literature searches.

## Data sharing

All sequencing data used in the present study is publicly available as part of the open-access MalariaGEN Plasmodium falciparum Community Project v7 (Pf7)^13^

## ACKNOWLEDGMENTS

This study was funded by the Bill & Melinda Gates Foundation (grant numbers OPP11188166 and OPP1204268) and by The Global Fund to Fight AIDS, Tuberculosis and Malaria (grant number 20864-007-44). This publication uses open-access data from the MalariaGEN *Plasmodium falciparum* Community Project v7 (Pf7) as described in https://doi.org/10.12688/wellcomeopenres.18681.1. The authors wish to thank all the patients and guardians who generously agreed to provide blood samples. We are indebted to all researchers who contributed samples to the Community Project since its inception, many of which were analyzed in the present work.

## AUTHOR CONTRIBUTION

AAN, LAE, MMAH, IA, EA, TA, GAA, PB, GIB, MBA, AC, DJC, UD, MD, AD, AMD, PD, RF, CF, AG, DI, ML, OMA, SA, ARU organized or carried out sample collections. SG conducted laboratory analyses. JA, RP, SG, CA produced genomic data. OM, VW, NW, ZB, WH performed data analyses. OM, VS, DPK designed and coordinated the project. OM, ZB drafted the manuscript.

## REFERENCES

1. White NJ, Pukrittayakamee S, Hien TT, Faiz MA, Mokuolu OA, Dondorp AM. Malaria. Lancet 2014; 383(9918): 723–35.

2. World Health Organization. World Malaria Report 2022. Geneva: World Health Organization, 2022.

3. Ekland EH, Fidock DA. Advances in understanding the genetic basis of antimalarial drug resistance. Curr Opin Microbiol 2007; 10: 363–70.

4. Manske M, Miotto O, Campino S, et al. Analysis of Plasmodium falciparum diversity in natural infections by deep sequencing. Nature 2012; 487(7407): 375–9.

5. Imwong M, Suwannasin K, Kunasol C, et al. The spread of artemisinin-resistant Plasmodium falciparum in the Greater Mekong subregion: a molecular epidemiology observational study. Lancet Infect Dis 2017; 17(5): 491–7.

6. Hamilton WL, Amato R, van der Pluijm RW, et al. Evolution and expansion of multidrugresistant malaria in southeast Asia: a genomic epidemiology study. Lancet Infect Dis 2019; 19(9): 943–51.

7. Wasakul V, Disratthakit A, Mayxay M, et al. Malaria outbreak in Laos driven by a selective sweep for Plasmodium falciparum kelch13 R539T mutants: a genetic epidemiology analysis. Lancet Infect Dis 2022.

8. MalariaGEN, Ahouidi A, Ali M, et al. An open dataset of Plasmodium falciparum genome variation in 7,000 worldwide samples. Wellcome Open Res 2021; 6: 42.

9. Miotto O, Almagro-Garcia J, Manske M, et al. Multiple populations of artemisinin-resistant Plasmodium falciparum in Cambodia. Nature Genetics 2013; 45(6): 648–55.

10. Miotto O, Amato R, Ashley EA, et al. Genetic architecture of artemisinin-resistant Plasmodium falciparum. Nat Genet 2015; 47(3): 226–34.

11. Amambua-Ngwa A, Amenga-Etego L, Kamau E, et al. Major subpopulations of Plasmodium falciparum in sub-Saharan Africa. Science 2019; 365(6455): 813–6.

12. Amambua-Ngwa A, Jeffries D, Amato R, et al. Consistent signatures of selection from genomic analysis of pairs of temporal and spatial Plasmodium falciparum populations from The Gambia. Sci Rep 2018; 8(1): 9687.

13. MalariaGEN. Pf7: an open dataset of Plasmodium falciparum genome variation in 20,000 worldwide samples. Wellcome Open Research 2022.

14. MalariaGEN Plasmodium falciparum Community Project. Genomic epidemiology of artemisinin resistant malaria. Elife 2016; 5.

15. Hill WG, Robertson A. Linkage disequilibrium in finite populations. Theoret Appl Genet 1968; 38: 226–31.

16. Singh S, Soe S, Weisman S, Barnwell JW, Perignon JL, Druilhe P. A conserved multi-gene family induces cross-reactive antibodies effective in defense against Plasmodium falciparum. PLoS One 2009; 4(4): e5410.

17. Otto TD, Bohme U, Sanders M, et al. Long read assemblies of geographically dispersed Plasmodium falciparum isolates reveal highly structured subtelomeres. Wellcome Open Res 2018; 3: 52.

18. Letcher B, Maciuca S, Iqbal Z. Gene conversion drives allelic dimorphism in two paralogous surface antigens of the malaria parasite <em>P. falciparum</em>. bioRxiv 2023: 2023.02.27.530215.

19. Tanabe K, Mackay M, Goman M, Scaife JG. Allelic dimorphism in a surface antigen gene of the malaria parasite Plasmodium falciparum. J Mol Biol 1987; 195(2): 273–87.

20. Roy SW, Ferreira MU, Hartl DL. Evolution of allelic dimorphism in malarial surface antigens. Heredity 2008; 100(2): 103–10.

21. Ferreira MU, Kaneko O, Kimura M, Liu Q, Kawamoto F, Tanabe K. Allelic diversity at the merozoite surface protein-1 (MSP-1) locus in natural Plasmodium falciparum populations: a brief overview. Memorias do Instituto Oswaldo Cruz 1998; 93(5): 631–8.

22. Miller LH, Roberts T, Shahabuddin M, McCutchan TF. Analysis of sequence diversity in the Plasmodium falciparum merozoite surface protein-1 (MSP-1). Molecular and biochemical parasitology 1993; 59(1): 1–14.

23. Kadekoppala M, Holder AA. Merozoite surface proteins of the malaria parasite: the MSP1 complex and the MSP7 family. International journal for parasitology 2010; 40(10): 1155–61.

24. Lin CS, Uboldi AD, Marapana D, et al. The merozoite surface protein 1 complex is a platform for binding to human erythrocytes by Plasmodium falciparum. J Biol Chem 2014; 289(37): 25655–69.

25. Lin CS, Uboldi AD, Epp C, et al. Multiple Plasmodium falciparum Merozoite Surface Protein 1 Complexes Mediate Merozoite Binding to Human Erythrocytes. J Biol Chem 2016; 291(14): 7703–15.

26. Das S, Hertrich N, Perrin AJ, et al. Processing of Plasmodium falciparum Merozoite Surface Protein MSP1 Activates a Spectrin-Binding Function Enabling Parasite Egress from RBCs. Cell Host Microbe 2015; 18(4): 433–44.

27. Koussis K, Withers-Martinez C, Yeoh S, et al. A multifunctional serine protease primes the malaria parasite for red blood cell invasion. EMBO J 2009; 28(6): 725–35.

28. Arisue N, Palacpac NMQ, Tougan T, Horii T. Characteristic features of the SERA multigene family in the malaria parasite. Parasit Vectors 2020; 13(1): 170.

29. Ruecker A, Shea M, Hackett F, et al. Proteolytic activation of the essential parasitophorous vacuole cysteine protease SERA6 accompanies malaria parasite egress from its host erythrocyte. J Biol Chem 2012; 287(45): 37949–63.

30. Sargeant TJ, Marti M, Caler E, et al. Lineage-specific expansion of proteins exported to erythrocytes in malaria parasites. Genome Biol 2006; 7(2): R12.

31. Vincensini L, Richert S, Blisnick T, et al. Proteomic analysis identifies novel proteins of the Maurer’s clefts, a secretory compartment delivering Plasmodium falciparum proteins to the surface of its host cell. Mol Cell Proteomics 2005; 4(4): 582–93.

32. Spycher C, Rug M, Pachlatko E, et al. The Maurer’s cleft protein MAHRP1 is essential for trafficking of PfEMP1 to the surface of Plasmodium falciparum-infected erythrocytes. Molecular Microbiology 2008; 68(5): 1300–14.

33. Nilsson S, Angeletti D, Wahlgren M, Chen Q, Moll K. Plasmodium falciparum antigen 332 is a resident peripheral membrane protein of Maurer’s clefts. PLoS One 2012; 7(11): e46980.

34. Spielmann T, Hawthorne PL, Dixon MW, et al. A cluster of ring stage-specific genes linked to a locus implicated in cytoadherence in Plasmodium falciparum codes for PEXEL-negative and PEXEL-positive proteins exported into the host cell. Mol Biol Cell 2006; 17(8): 3613–24.

35. Dixon MW, Kenny S, McMillan PJ, et al. Genetic ablation of a Maurer’s cleft protein prevents assembly of the Plasmodium falciparum virulence complex. Mol Microbiol 2011; 81(4): 982–93.

36. Nunes MC, Okada M, Scheidig-Benatar C, Cooke BM, Scherf A. Plasmodium falciparum FIKK kinase members target distinct components of the erythrocyte membrane. PLoS One 2010; 5(7): e11747.

37. Gardiner DL, Spielmann T, Dixon MW, et al. CLAG 9 is located in the rhoptries of Plasmodium falciparum. Parasitol Res 2004; 93(1): 64–7.

38. Kelleher J, Wong Y, Wohns AW, Fadil C, Albers PK, McVean G. Inferring whole-genome histories in large population datasets. Nat Genet 2019; 51(9): 1330–8.

39. Taylor SM, Cerami C, Fairhurst RM. Hemoglobinopathies: slicing the Gordian knot of Plasmodium falciparum malaria pathogenesis. PLoS Pathog 2013; 9(5): e1003327.

40. Cyrklaff M, Sanchez CP, Kilian N, et al. Hemoglobins S and C interfere with actin remodeling in Plasmodium falciparum-infected erythrocytes. Science 2011; 334(6060): 1283–6.

41. Paradis E, Schliep K. ape 5.0: an environment for modern phylogenetics and evolutionary analyses in R. Bioinformatics 2019; 35(3): 526–8.

42. Krzywinski M, Schein J, Birol I, et al. Circos: an information aesthetic for comparative genomics. Genome Res 2009; 19(9): 1639–45.

43. Schaffner SF, Taylor AR, Wong W, Wirth DF, Neafsey DE. hmmIBD: software to infer pairwise identity by descent between haploid genotypes. Malar J 2018; 17(1): 196.

44. Iqbal Z, Caccamo M, Turner I, Flicek P, McVean G. De novo assembly and genotyping of variants using colored de Bruijn graphs. Nature Genetics 2012; 44(2): 226–32.

45. Manske HM, Kwiatkowski DP. LookSeq: a browser-based viewer for deep sequencing data. Genome research 2009; 19(11): 2125–32.

