## Supplementary Tables and Figures for "A complex *Plasmodium falciparum* cryptotype circulating at low frequency across the African continent"

Miotto O *et al*.

#### SUPPLEMENTARY MATERIALS

#### SUPPLEMENTARY TABLES

##### Supplementary Table 1 – Highly Differentiated non-synonymous coding SNPs in AF1.

Each row represents one non-synonymous coding SNP (n=175) that exhibits mean $F_{ST}\geq0.5$ between AF1 and the three African macro-regions. The columns show: chromosome number and position of the SNP within the chromosome; mean $F_{ST}$; ID and description of the gene containing the SNP; the amino acid mutation caused by the non-reference allele; and the estimated frequencies of the non-reference allele in AF1 and the following populations: West Africa (WAF), Central Africa (CAF), East Africa (EAF), South Asia (SAS), Western Southeast Asia (WSEA), Eastern Southeast Asia (ESEA), Oceania (OCE) and South America (SAM). SNPs with $F_{ST}\geq0.75$ are highlighted in **bold** type. SNPs that had high missingness only in sWGA-processed samples (see main text and Methods) are indicated by a coloured background in their Chr/Pos fields. Reported positions, identifiers, mutations and non-reference alleles are all with respect to the 3D7 V3 reference genome. To aid visualization, the backgrounds of the $F_{ST}$ and frequency estimates were coloured so that higher values are represented by more saturated background colours.

| Chr | Pos | $F_{ST}$ | Gene ID | Gene Description | Mutation | AF1 | WAF | CAF | EAF | SAS | WSEA | ESEA | OCE | SAM |
|---|---|---|---|---|---|---|---|---|---|---|---|---|---|---|
| 1 | 114559 | 0.62 | PF3D7_0102500 | erythrocyte binding antigen-181 | N414I | 0.82 | 0.03 | 0.02 | 0.09 | 0.72 | 0.99 | 0.99 | 0.78 | 0.08 |
| 1 | 114724 | 0.66 | | | R359K | 0.81 | 0.01 | 0 | 0.02 | 0.03 | 0.34 | 0.31 | 0.59 | 0.08 |
| 1 | 132574 | 0.51 | PF3D7_0103000 | vacuolar protein sorting-associated protein VTA1, putative | E182K | 0.81 | 0.10 | 0.09 | 0.09 | 0.02 | 0 | 0 | 0 | 0 |
| 1 | 180034 | 0.63 | PF3D7_0104100 | protein E140, putative | E540D | 0.09 | 0.87 | 0.89 | 0.88 | 0.86 | 0.85 | 0.87 | 0.88 | 0.47 |
| 1 | 527210 | 0.56 | PF3D7_0113800 | DBL containing protein, unknown function | N35S | 0.83 | 0.10 | 0.06 | 0.09 | 0.07 | 0.02 | 0 | 0.08 | 0.01 |
| **1** | **563776** | **0.80** | **PF3D7_0114700** | **PIR protein** | **A300V** | **0.93** | **0.04** | **0.05** | **0.03** | **0.01** | **0** | **0** | **0** | **0.04** |
| 2 | 153549 | 0.65 | PF3D7_0203100 | protein kinase, putative | E1145K | 0.80 | 0.01 | 0.01 | 0.01 | 0 | 0 | 0 | 0 | 0 |
| **2** | **301238** | **0.89** | **PF3D7_0207500** | **serine repeat antigen 6** | **Q290K** | **0.95** | **0** | **0** | **0.01** | **0** | **0** | **0** | **0** | **0** |
| **2** | **303754** | **0.90** | **PF3D7_0207600** | **serine repeat antigen 5** | **K945E** | **0.95** | **0** | **0** | **0.01** | **0** | **0** | **0** | **0** | **0** |
| **2** | **303786** | **0.90** | | | **R934H** | **0.95** | **0** | **0** | **0.01** | **0** | **0** | **0** | **0** | **0** |
| **2** | **305718** | **0.88** | | | **I330L** | **0.95** | **0** | **0** | **0.04** | **0.75** | **1.00** | **0.99** | **0.93** | **1.00** |
| **2** | **306406** | **0.84** | | | **K159E** | **0.93** | **0.01** | **0** | **0.03** | **0.79** | **1.00** | **1.00** | **0.92** | **1.00** |
| 2 | 315716 | 0.59 | PF3D7_0207800 | serine repeat antigen 3 | T234P | 0.05 | 0.83 | 0.81 | 0.78 | 0.89 | 1.00 | 0.99 | 1.00 | 0.91 |
| 2 | 320853 | 0.67 | PF3D7_0207900 | serine repeat antigen 2 | P89S | 1.00 | 0.24 | 0.14 | 0.22 | 0 | 0 | 0 | 0.02 | 0 |
| 2 | 337669 | 0.56 | PF3D7_0208300 | conserved Plasmodium protein, unknown function | Y38N | 0.75 | 0.03 | 0.01 | 0.03 | 0.01 | 0 | 0 | 0 | 0 |
| 2 | 373697 | 0.52 | PF3D7_0209000 | transmission-blocking target antigen s230 | S1087Y | 0.86 | 0.25 | 0.14 | 0.05 | 0.28 | 0.17 | 0.08 | 0 | 0.10 |
| 2 | 735613 | 0.62 | PF3D7_0217900 | thioesterase/thiol ester dehydrase-isomerase, putative | S462T | 0.81 | 0.06 | 0.01 | 0.03 | 0 | 0 | 0 | 0 | 0 |
| **2** | **784067** | **0.79** | **PF3D7_0219700** | **gametocyte exported protein 20** | **Y182H** | **0.89** | **0.01** | **0.01** | **0.01** | **0** | **0** | **0** | **0** | **0.15** |
| 2 | 784379 | 0.65 | | | H78N | 0.92 | 0.16 | 0.10 | 0.08 | 0.15 | 0.07 | 0.02 | 0 | 0.15 |
| 2 | 814192 | 0.56 | PF3D7_0220300 | Plasmodium exported protein, unknown function | P92A | 0.72 | 0 | 0 | 0.01 | 0 | 0 | 0 | 0 | 0 |
| 3 | 865666 | 0.63 | PF3D7_0320700 | signal peptidase complex subunit 2 | M78L | 0.89 | 0.08 | 0.09 | 0.11 | 0 | 0 | 0 | 0 | 0.48 |
| **4** | **103881** | **0.85** | **PF3D7_0401800** | **Plasmodium exported protein (PHISTb), unknown function** | **K515R** | **0.93** | **0** | **0.01** | **0.02** | **0.41** | **0.84** | **0.68** | **0.71** | **0** |
| 4 | 103987 | 0.53 | | | K480E | 0.04 | 0.73 | 0.72 | 0.80 | 0.77 | 0.89 | 0.57 | 0.97 | 0.69 |

| | | | | | | | | | | | | | | |
|---|---|---|---|---|---|---|---|---|---|---|---|---|---|---|
| 4 | 104157 | 0.58 | | | H423P | 0.91 | 0.09 | 0.20 | 0.17 | 0.21 | 0.13 | 0.33 | 0.14 | 0 |
| 4 | 110821 | 0.56 | PF3D7_0401900 | acyl-CoA synthetase | I159L | 0.98 | 0.20 | 0.30 | 0.25 | 0.59 | 0.82 | 0.47 | 0.70 | 0.11 |
| **4** | **464779** | **0.79** | **PF3D7_0410000** | **erythrocyte vesicle protein 1** | **D819Y** | **0.90** | **0.02** | **0** | **0.03** | **0** | **0** | **0** | **0** | **0.05** |
| 4 | 1103709 | 0.58 | PF3D7_0424400 | surface-associated interspersed protein 4.2 (SURFIN 4.2) | W1247. | 0.76 | 0.03 | 0.03 | 0 | 0 | 0 | 0 | 0 | 0 |
| 4 | 1113576 | 0.61 | PF3D7_0424600 | Plasmodium exported protein (PHISTb) | K233N | 0.89 | 0.06 | 0.07 | 0.21 | 0.68 | 0.69 | 0.51 | 0.95 | 0.51 |
| 6 | 851783 | 0.58 | PF3D7_0620400 | merozoite surface protein 10 | K391N | 0.89 | 0.09 | 0.09 | 0.23 | 0.10 | 0.02 | 0.01 | 0 | 0.43 |
| 7 | 712688 | 0.57 | PF3D7_0716200 | PDCD2 domain-containing protein, putative | G74R | 0.74 | 0.02 | 0 | 0.01 | 0 | 0 | 0 | 0 | 0 |
| 7 | 1359488 | 0.65 | PF3D7_0731500 | erythrocyte binding antigen-175 | K478N | 0.90 | 0.05 | 0.08 | 0.17 | 0.07 | 0.07 | 0.22 | 0.38 | 0.08 |
| 8 | 1056829 | 0.67 | PF3D7_0824200 | conserved Plasmodium protein, unknown function | L474I | 0.07 | 0.93 | 0.93 | 0.78 | 0.71 | 0.63 | 0.42 | 0.62 | 0.88 |
| 8 | 1238850 | 0.60 | PF3D7_0828800 | GPI-anchored micronemal antigen | V218I | 0.79 | 0.03 | 0.02 | 0.03 | 0 | 0 | 0 | 0 | 0 |
| 8 | 1296885 | 0.59 | PF3D7_0830500 | tryptophan-rich antigen | F426Y | 0.82 | 0.05 | 0.04 | 0.09 | 0.60 | 0.83 | 0.83 | 0.77 | 0.26 |
| 8 | 1311901 | 0.69 | PF3D7_0830800 | surface-associated interspersed protein 8.2 (SURFIN 8.2) | P422R | 0.95 | 0.14 | 0.10 | 0.13 | 0.13 | 0.09 | 0.13 | 0.06 | 0.03 |
| 8 | 1311927 | 0.51 | | | N431H | 0.98 | 0.27 | 0.29 | 0.29 | 0.45 | 0.56 | 0.57 | 0.32 | 0.53 |
| 8 | 1311929 | 0.63 | | | N431K | 0.95 | 0.15 | 0.17 | 0.17 | 0.25 | 0.26 | 0.25 | 0.22 | 0.20 |
| 8 | 1311938 | 0.61 | | | F434L | 0.95 | 0.16 | 0.18 | 0.19 | 0.27 | 0.26 | 0.25 | 0.22 | 0.20 |
| 8 | 1311959 | 0.52 | | | L441F | 0.95 | 0.23 | 0.22 | 0.28 | 0.30 | 0.41 | 0.32 | 0.32 | 0.21 |
| 8 | 1312185 | 0.62 | | | N517H | 0.95 | 0.13 | 0.19 | 0.19 | 0.11 | 0.24 | 0.17 | 0.11 | 0.03 |
| 8 | 1344521 | 0.66 | PF3D7_0831400 | Plasmodium exported protein, unknown function | N265D | 0.98 | 0.46 | 0.05 | 0.09 | 0.63 | 0.82 | 0.96 | 0.67 | 0.50 |
| 8 | 1344529 | 0.66 | | | I262K | 0.98 | 0.45 | 0.05 | 0.09 | 0.62 | 0.82 | 0.96 | 0.67 | 0.50 |
| **9** | **82156** | **0.76** | **PF3D7_0901700** | **Plasmodium exported protein (hyp5), unknown function** | **Y177C** | **0.91** | **0.01** | **0.02** | **0.09** | **0.08** | **0.18** | **0.28** | **0.25** | **0.40** |
| 9 | 82238 | 0.63 | | | N150D | 0.93 | 0.08 | 0.13 | 0.23 | 0.21 | 0.37 | 0.36 | 0.36 | 0.40 |
| 9 | 84790 | 0.65 | PF3D7_0901800 | Plasmodium exported protein, unknown function | F56S | 0.91 | 0.07 | 0.12 | 0.12 | 0.11 | 0.15 | 0.35 | 0 | 0.37 |
| 9 | 85450 | 0.50 | | | T217N | 0.91 | 0.13 | 0.19 | 0.33 | 0.41 | 0.72 | 0.78 | 0.66 | 0.53 |
| **9** | **465933** | **0.79** | **PF3D7_0910200** | **conserved Plasmodium protein, unknown function** | **T459A** | **0.91** | **0.02** | **0.02** | **0.04** | **0** | **0** | **0** | **0** | **0** |
| 9 | 527158 | 0.53 | PF3D7_0911500 | conserved Plasmodium protein, unknown function | C147F | 0.81 | 0.07 | 0.09 | 0.10 | 0.02 | 0 | 0 | 0 | 0 |
| 9 | 778894 | 0.67 | PF3D7_0918900 | gamma-glutamylcysteine synthetase | N446S | 0.84 | 0.03 | 0.02 | 0.02 | 0 | 0 | 0 | 0 | 0 |
| **9** | **781952** | **0.75** | **PF3D7_0919000** | **nucleosome assembly protein** | **I76V** | **0.87** | **0.01** | **0.01** | **0.01** | **0.01** | **0.06** | **0.05** | **0.36** | **0** |
| 9 | 799189 | 0.66 | PF3D7_0919500 | major facilitator superfamily domain-containing protein, put. | E231V | 0.87 | 0.07 | 0.06 | 0.04 | 0 | 0 | 0 | 0 | 0 |
| **9** | **1175905** | **0.83** | **PF3D7_0929400** | **high molecular weight rhoptry protein 2** | **A235T** | **0.93** | **0.02** | **0.02** | **0.03** | **0** | **0** | **0** | **0** | **0.06** |
| **9** | **1202267** | **0.89** | **PF3D7_0930300** | **merozoite surface protein 1** | **F152L** | **0.95** | **0.01** | **0** | **0.02** | **0.04** | **0.27** | **0.11** | **0.15** | **0.10** |
| **9** | **1202292** | **0.85** | | | **E161Q** | **0.93** | **0.01** | **0** | **0.02** | **0.04** | **0.27** | **0.11** | **0.15** | **0.10** |
| **9** | **1202596** | **0.92** | | | **T262K** | **0.97** | **0.01** | **0.01** | **0.03** | **0.04** | **0.33** | **0.11** | **0.04** | **0.10** |
| **9** | **1202605** | **0.92** | | | **A265E** | **0.97** | **0.01** | **0.01** | **0.03** | **0.04** | **0.33** | **0.11** | **0.04** | **0.10** |
| **9** | **1202649** | **0.92** | | | **Q280K** | **0.97** | **0.01** | **0.01** | **0.03** | **0.04** | **0.27** | **0.11** | **0.05** | **0.10** |
| **9** | **1202664** | **0.83** | | | **D285N** | **0.97** | **0.04** | **0.07** | **0.07** | **0.08** | **0.27** | **0.11** | **0.05** | **0.31** |
| **9** | **1202665** | **0.83** | | | **D285A** | **0.97** | **0.04** | **0.07** | **0.07** | **0.08** | **0.27** | **0.11** | **0.05** | **0.31** |
| **9** | **1202669** | **0.83** | | | **N286K** | **0.97** | **0.04** | **0.07** | **0.07** | **0.08** | **0.27** | **0.11** | **0.05** | **0.31** |
| **9** | **1202913** | **0.93** | | | **K368E** | **0.97** | **0.01** | **0** | **0.02** | **0.13** | **0.29** | **0.13** | **0.15** | **0.10** |

| | | | | | | | | | | | | | | |
|---|---|---|---|---|---|---|---|---|---|---|---|---|---|---|
| 9 | 1203652 | 0.61 | | | L614R | 1.00 | 0.26 | 0.34 | 0.14 | 0.07 | 0.05 | 0.04 | 0.19 | 0.02 |
| **9** | **1203952** | **0.96** | | | **S714N** | **1.00** | **0.03** | **0.01** | **0.02** | **0** | **0** | **0** | **0** | **0** |
| **9** | **1205118** | **0.93** | | | **H1103N** | **0.97** | **0.01** | **0** | **0.02** | **0.04** | **0.29** | **0.11** | **0.16** | **0.24** |
| **9** | **1205120** | **0.93** | | | **H1103Q** | **0.97** | **0.01** | **0** | **0.02** | **0.04** | **0.29** | **0.11** | **0.16** | **0.10** |
| **9** | **1205121** | **0.93** | | | **N1104H** | **0.97** | **0.01** | **0** | **0.02** | **0.04** | **0.29** | **0.11** | **0.16** | **0.10** |
| **9** | **1205151** | **0.91** | | | **N1114Y** | **0.96** | **0.01** | **0** | **0** | **0** | **0** | **0** | **0** | **0.10** |
| **9** | **1205284** | **0.93** | | | **V1158E** | **0.97** | **0.01** | **0** | **0.02** | **0.04** | **0.28** | **0.10** | **0.14** | **0.10** |
| **9** | **1205314** | **0.93** | | | **N1168S** | **0.97** | **0.01** | **0** | **0.02** | **0.04** | **0.29** | **0.11** | **0.14** | **0.10** |
| **9** | **1205324** | **0.93** | | | **K1171N** | **0.97** | **0.01** | **0** | **0.02** | **0.04** | **0.29** | **0.10** | **0.14** | **0.10** |
| **9** | **1205329** | **0.93** | | | **R1173K** | **0.97** | **0.01** | **0** | **0.02** | **0.04** | **0.29** | **0.10** | **0.14** | **0.10** |
| **9** | **1205343** | **0.93** | | | **I1178L** | **0.97** | **0.01** | **0** | **0.02** | **0.04** | **0.29** | **0.10** | **0.14** | **0.10** |
| **9** | **1205355** | **0.93** | | | **L1182F** | **0.97** | **0.01** | **0** | **0.02** | **0.04** | **0.28** | **0.10** | **0.13** | **0.10** |
| **9** | **1205360** | **0.93** | | | **N1183K** | **0.97** | **0.01** | **0** | **0.02** | **0.04** | **0.29** | **0.10** | **0.13** | **0.10** |
| **9** | **1205370** | **0.93** | | | **H1187N** | **0.97** | **0.01** | **0** | **0.02** | **0.04** | **0.29** | **0.10** | **0.14** | **0.10** |
| **9** | **1205377** | **0.93** | | | **G1189E** | **0.97** | **0.01** | **0** | **0.02** | **0.04** | **0.29** | **0.10** | **0.14** | **0.10** |
| **9** | **1205395** | **0.93** | | | **F1195Y** | **0.97** | **0.01** | **0** | **0.02** | **0.04** | **0.28** | **0.10** | **0.14** | **0.10** |
| **9** | **1205424** | **0.93** | | | **T1205A** | **0.97** | **0.01** | **0** | **0.02** | **0.04** | **0.29** | **0.11** | **0.14** | **0.10** |
| 9 | 1316936 | 0.52 | PF3D7_0933100 | conserved Plasmodium protein, unknown function | V606A | 0.91 | 0.12 | 0.19 | 0.28 | 0.05 | 0 | 0 | 0.01 | 0.95 |
| **9** | **1417854** | **0.79** | **PF3D7_0935800** | **cytoadherence linked asexual protein 9** | **T779S** | **0.97** | **0.15** | **0.06** | **0.05** | **0** | **0** | **0** | **0** | **0** |
| 9 | 1419023 | 0.74 | | | K1098Q | 0.97 | 0.12 | 0.13 | 0.11 | 0.32 | 0.21 | 0.31 | 0.50 | 0.87 |
| 9 | 1420566 | 0.67 | PF3D7_0935900 | ring-exported protein 1 | E687Q | 0.84 | 0.03 | 0.04 | 0.01 | 0 | 0 | 0 | 0 | 0.04 |
| **9** | **1427697** | **0.89** | | | **S77.** | **0.98** | **0.06** | **0.02** | **0.02** | **0** | **0** | **0** | **0** | **0.32** |
| **9** | **1427982** | **0.87** | **PF3D7_0936000** | **ring-exported protein 2** | **E14A** | **0.98** | **0.06** | **0.02** | **0.05** | **0.02** | **0.31** | **0.07** | **0.10** | **0.86** |
| 9 | 1428013 | 0.69 | | | Y4N | 0.98 | 0.11 | 0.13 | 0.23 | 0.47 | 0.68 | 0.44 | 0.36 | 0.88 |
| 10 | 326012 | 0.50 | PF3D7_1008000 | histone deacetylase 2 | N1668D | 0.84 | 0.14 | 0.08 | 0.18 | 0.04 | 0 | 0 | 0.03 | 0 |
| 10 | 354221 | 0.59 | PF3D7_1008500 | protein GPR89, putative | L840P | 0.96 | 0.18 | 0.18 | 0.25 | 0.19 | 0.06 | 0.01 | 0 | 0 |
| 10 | 559189 | 0.50 | PF3D7_1014100 | merozoite surface protein MSA180 | N1344H | 0.71 | 0.06 | 0.02 | 0 | 0 | 0 | 0 | 0 | 0 |
| 10 | 562016 | 0.52 | PF3D7_1014200 | male gamete fusion factor HAP2, putative | D445N | 0.85 | 0.29 | 0.09 | 0.05 | 0.11 | 0.07 | 0.02 | 0.02 | 0.04 |
| 10 | 563297 | 0.51 | | | H872Y | 0.74 | 0.09 | 0.02 | 0.02 | 0 | 0 | 0 | 0 | 0 |
| 10 | 571802 | 0.57 | PF3D7_1014300 | SPRY domain-containing protein, putative | H2000Y | 0.86 | 0.15 | 0.08 | 0.09 | 0.01 | 0.01 | 0.02 | 0 | 0 |
| 10 | 578813 | 0.52 | PF3D7_1014500 | conserved Plasmodium protein, unknown function | I745M | 0.74 | 0.07 | 0.02 | 0.02 | 0.01 | 0 | 0 | 0 | 0 |
| 10 | 582046 | 0.51 | PF3D7_1014600 | transcriptional coactivator ADA2 | D2349Y | 0.72 | 0.06 | 0.01 | 0.01 | 0.01 | 0 | 0 | 0 | 0 |
| 10 | 653563 | 0.51 | PF3D7_1016300 | glycophorin binding protein | R2Q | 0.71 | 0.03 | 0.04 | 0.01 | 0 | 0 | 0 | 0 | 0 |
| 10 | 1038679 | 0.63 | PF3D7_1024800 | exported protein 3 | Q1332L | 0.88 | 0.10 | 0.10 | 0.07 | 0 | 0 | 0.01 | 0 | 0.25 |
| 10 | 1285388 | 0.50 | PF3D7_1031900 | conserved Plasmodium protein, unknown function | F417L | 1.00 | 0.38 | 0.30 | 0.31 | 0.20 | 0.05 | 0.01 | 0.06 | 0.05 |
| 10 | 1325994 | 0.66 | PF3D7_1033100 | S-adenosylmethionine decarboxylase/ornithine decarboxylase | N815Y | 0.98 | 0.36 | 0.11 | 0.09 | 0.03 | 0 | 0 | 0 | 0 |
| 10 | 1325996 | 0.66 | | | E814G | 0.98 | 0.36 | 0.11 | 0.09 | 0.03 | 0 | 0 | 0 | 0 |
| **10** | **1371865** | **0.88** | **PF3D7_1034500** | **armadillo repeat protein, putative** | **T336I** | **0.98** | **0.03** | **0.03** | **0.05** | **0.05** | **0.04** | **0** | **0** | **0** |

| | | | | | | | | | | | | | | |
|---|---|---|---|---|---|---|---|---|---|---|---|---|---|---|
| 10 | 1373309 | 0.54 | | | I817M | 0.97 | 0.27 | 0.24 | 0.25 | 0.25 | 0.18 | 0.36 | 0.05 | 0.04 |
| 10 | 1391865 | 0.50 | | | S141G | 0.02 | 0.69 | 0.69 | 0.74 | 0.73 | 0.86 | 0.85 | 0.72 | 0.12 |
| 10 | 1391943 | 0.58 | | | N167D | 0.02 | 0.76 | 0.77 | 0.78 | 0.84 | 0.91 | 0.93 | 0.88 | 1.00 |
| 10 | 1391973 | 0.57 | **PF3D7_1035100** | **probable protein, unknown function** | N177Y | 0.02 | 0.75 | 0.76 | 0.78 | 0.84 | 0.91 | 0.93 | 0.86 | 1.00 |
| **10** | **1392014** | **0.76** | | | **H190Q** | **0.03** | **0.88** | **0.92** | **0.90** | **0.90** | **0.98** | **0.94** | **0.90** | **1.00** |
| 10 | 1392155 | 0.55 | | | N237K | 0.78 | 0.04 | 0.04 | 0.07 | 0.04 | 0.07 | 0.08 | 0.35 | 0.83 |
| 10 | 1399580 | 0.52 | | | D129G | 0 | 0.68 | 0.75 | 0.61 | 0.15 | 0.09 | 0.09 | 0.06 | 0.01 |
| **10** | **1399594** | **0.95** | | | **S134T** | **1.00** | **0.02** | **0.02** | **0.03** | **0.04** | **0.03** | **0.04** | **0.01** | **0.89** |
| **10** | **1399634** | **0.92** | **PF3D7_1035300** | **glutamate-rich protein GLURP** | **V147G** | **1.00** | **0.04** | **0.03** | **0.04** | **0.04** | **0.03** | **0.04** | **0.06** | **0.89** |
| **10** | **1399636** | **0.91** | | | **Q148E** | **1.00** | **0.04** | **0.03** | **0.06** | **0.04** | **0.03** | **0.04** | **0.11** | **0.89** |
| **10** | **1399656** | **0.95** | | | **L154F** | **1.00** | **0.02** | **0.02** | **0.03** | **0.04** | **0.03** | **0.04** | **0.01** | **0.89** |
| **10** | **1399681** | **0.94** | | | **S163P** | **1.00** | **0.02** | **0.03** | **0.03** | **0.04** | **0.03** | **0.04** | **0.01** | **0.89** |
| 10 | 1404580 | 0.66 | PF3D7_1035400 | merozoite surface protein 3 | A129V | 0.94 | 0.11 | 0.11 | 0.17 | 0.23 | 0.28 | 0.31 | 0.21 | 0.10 |
| 10 | 1404591 | 0.57 | | | V133F | 0.99 | 0.24 | 0.25 | 0.27 | 0.41 | 0.44 | 0.45 | 0.52 | 0.10 |
| **10** | **1413597** | **0.79** | | | **G133D** | **0.88** | **0** | **0** | **0** | **0** | **0** | **0** | **0** | **0** |
| **10** | **1413618** | **0.83** | | | **N140S** | **0.93** | **0** | **0.03** | **0.02** | **0** | **0** | **0** | **0** | **0** |
| 10 | 1413659 | 0.62 | | | K154Q | 0.96 | 0.21 | 0.19 | 0.15 | 0.08 | 0.15 | 0.09 | 0.11 | 0.73 |
| 10 | 1413669 | 0.54 | **PF3D7_1035700** | **duffy binding-like merozoite surface protein** | L157. | 0.91 | 0.20 | 0.18 | 0.15 | 0.07 | 0.15 | 0.08 | 0.11 | 0.72 |
| 10 | 1413686 | 0.73 | | | N163D | 0.91 | 0.08 | 0.07 | 0.02 | 0.02 | 0.06 | 0.01 | 0.02 | 0.72 |
| **10** | **1414634** | **0.91** | | | **L479I** | **0.97** | **0** | **0.02** | **0.02** | **0** | **0** | **0** | **0** | **0** |
| 10 | 1415066 | 0.58 | | | K623E | 0.97 | 0.18 | 0.23 | 0.25 | 0.50 | 0.53 | 0.47 | 0.18 | 0.13 |
| **10** | **1421410** | **0.86** | **PF3D7_1035800** | **probable protein, unknown function** | **G293D** | **0.97** | **0.01** | **0.05** | **0.07** | **0.30** | **0.28** | **0.26** | **0.62** | **0.86** |
| **10** | **1456571** | **0.94** | **PF3D7_1036900** | **conserved Plasmodium protein, unknown function** | **S479I** | **0.97** | **0** | **0** | **0.01** | **0** | **0** | **0** | **0** | **0** |
| **10** | **1468777** | **0.80** | **PF3D7_1037000** | **DNA polymerase zeta catalytic subunit, putative** | **N1968S** | **0.90** | **0.01** | **0.01** | **0.02** | **0** | **0** | **0** | **0** | **0** |
| **10** | **1483355** | **0.92** | **PF3D7_1037400** | **conserved Plasmodium protein, unknown function** | **E667Q** | **0.98** | **0.02** | **0.02** | **0.02** | **0** | **0** | **0** | **0** | **0** |
| 10 | 1485068 | 0.56 | | | S96T | 0.87 | 0.11 | 0.17 | 0.10 | 0.03 | 0 | 0 | 0.22 | 0.20 |
| **10** | **1503043** | **0.90** | **PF3D7_1037900** | **conserved Plasmodium protein, unknown function** | **N154S** | **0.97** | **0.03** | **0.04** | **0.01** | **0** | **0** | **0** | **0** | **0.07** |
| 10 | 1519152 | 0.61 | PF3D7_1038400 | gametocyte-specific protein | R44S | 0.76 | 0 | 0 | 0 | 0 | 0 | 0 | 0 | 0 |
| **10** | **1548519** | **0.88** | **PF3D7_1038500** | **Plasmodium exported protein, unknown function** | **F354C** | **0.98** | **0.05** | **0.02** | **0.04** | **0.11** | **0.07** | **0.13** | **0.32** | **0.09** |
| 10 | 1552485 | 0.55 | PF3D7_1038600 | Plasmodium exported protein, unknown function | Q190L | 0 | 0.79 | 0.70 | 0.62 | 0.32 | 0.09 | 0.07 | 0.80 | 0.35 |
| 10 | 1552843 | 0.53 | | | F71I | 0.70 | 0 | 0 | 0 | 0 | 0 | 0 | 0 | 0 |
| **10** | **1570402** | **0.78** | **PF3D7_1039000** | **serine/threonine protein kinase, FIKK family** | **E515A** | **0.89** | **0.02** | **0** | **0.02** | **0** | **0** | **0.06** | **0** | **0.01** |
| 11 | 120959 | 0.56 | PF3D7_1102500 | Plasmodium exported protein (PHISTb) | E110Q | 0.95 | 0.22 | 0.19 | 0.24 | 0.56 | 0.60 | 0.93 | 0.78 | 0.01 |
| 11 | 123946 | 0.52 | PF3D7_1102600 | gametocyte exported protein 14 | L230I | 0.75 | 0.04 | 0.04 | 0.04 | 0.03 | 0 | 0 | 0 | 0 |
| 11 | 123955 | 0.51 | | | L227F | 0.74 | 0.05 | 0.04 | 0.05 | 0.03 | 0 | 0 | 0 | 0 |
| 11 | 137245 | 0.52 | PF3D7_1102900 | Plasmodium exported protein (hyp11), unknown function | D151Y | 0.77 | 0.04 | 0.05 | 0.08 | 0 | 0 | 0 | 0 | 0 |
| 11 | 1484887 | 0.60 | PF3D7_1137900 | conserved Plasmodium protein, unknown function | H505D | 0.87 | 0.07 | 0.09 | 0.12 | 0.01 | 0 | 0 | 0.01 | 0.02 |
| 11 | 1640343 | 0.59 | PF3D7_1140900 | conserved Plasmodium protein, unknown function | G865E | 0.96 | 0.27 | 0.21 | 0.13 | 0.03 | 0.03 | 0 | 0 | 0.02 |

| | | | | | | | | | | | | | | |
|---|---|---|---|---|---|---|---|---|---|---|---|---|---|---|
| 11 | 1642013 | 0.60 | | | Y364F | 0.96 | 0.22 | 0.24 | 0.13 | 0.01 | 0.02 | 0 | 0 | 0.24 |
| 11 | 1664948 | 0.58 | PF3D7_1141400 | phosphatidylinositol N-acetylglucosaminyltransferase subunit H, putative | D254H | 0.85 | 0.13 | 0.09 | 0.07 | 0.02 | 0 | 0 | 0 | 0.02 |
| 11 | 1684025 | 0.56 | PF3D7_1142100 | conserved Plasmodium protein, unknown function | N2020K | 0.83 | 0.10 | 0.08 | 0.06 | 0.02 | 0.01 | 0 | 0 | 0 |
| 11 | 1862558 | 0.53 | PF3D7_1147000 | sporozoite asparagine-rich protein | D1600E | 0.70 | 0.01 | 0 | 0.01 | 0 | 0 | 0 | 0 | 0 |
| 11 | 1863226 | 0.53 | | | E1378K | 0.70 | 0.01 | 0.01 | 0.01 | 0.03 | 0 | 0 | 0 | 0 |
| 11 | 1941726 | 0.55 | PF3D7_1148800 | Plasmodium exported protein (hyp11), unknown function | T86I | 0.79 | 0.06 | 0.06 | 0.07 | 0.53 | 0.70 | 0.34 | 0.88 | 0.10 |
| 11 | 1956584 | 0.63 | | | G2047E | 0.79 | 0.01 | 0.02 | 0.01 | 0 | 0 | 0 | 0 | 0 |
| 11 | 1957091 | 0.63 | | | P2216Q | 0.78 | 0 | 0.01 | 0.01 | 0 | 0.01 | 0 | 0 | 0 |
| 11 | 1967527 | 0.57 | **PF3D7_1149000** | **antigen 332, DBL-like protein** | E5695K | 0.97 | 0.21 | 0.30 | 0.19 | 0.02 | 0.02 | 0.04 | 0.01 | 0.20 |
| **11** | **1967597** | **0.94** | | | **V5718E** | **0.97** | **0** | **0** | **0** | **0** | **0** | **0** | **0** | **0** |
| **11** | **1967951** | **0.86** | | | **S5836L** | **0.92** | **0** | **0** | **0** | **0** | **0** | **0** | **0** | **0** |
| **11** | **1976808** | **0.78** | | | **P93A** | **0.88** | **0** | **0** | **0** | **0** | **0** | **0** | **0** | **0** |
| 11 | 1977393 | 0.59 | | | Q288E | 0.74 | 0 | 0 | 0 | 0 | 0 | 0 | 0 | 0 |
| **11** | **1979012** | **0.84** | **PF3D7_1149200** | **ring-infected erythrocyte surface antigen** | **K827N** | **0.03** | **0.90** | **0.98** | **0.95** | **0.92** | **0.99** | **1.00** | **0.98** | **1.00** |
| 11 | 1979127 | 0.74 | | | I866L | 0.86 | 0.03 | 0.01 | 0 | 0.01 | 0 | 0 | 0 | 0 |
| 11 | 1979200 | 0.54 | | | A890V | 0.97 | 0.26 | 0.20 | 0.33 | 0.74 | 0.86 | 0.97 | 0.98 | 0.02 |
| **11** | **2001089** | **0.83** | | | **Y8N** | **0.96** | **0.07** | **0.05** | **0.03** | **0.06** | **0.04** | **0.01** | **0.11** | **0** |
| 11 | 2002527 | 0.66 | | | D453E | 0.06 | 0.87 | 0.85 | 0.87 | 0.87 | 0.77 | 0.92 | 0.93 | 0.71 |
| 11 | 2002901 | 0.67 | | | T578I | 0.06 | 0.89 | 0.86 | 0.87 | 0.92 | 0.89 | 0.96 | 0.93 | 1.00 |
| 11 | 2002933 | 0.58 | **PF3D7_1149600** | **DnaJ protein, putative** | V589M | 0.06 | 0.83 | 0.81 | 0.82 | 0.84 | 0.84 | 0.93 | 0.88 | 1.00 |
| 11 | 2002960 | 0.66 | | | H598Y | 0.06 | 0.89 | 0.86 | 0.86 | 0.90 | 0.88 | 0.95 | 0.94 | 1.00 |
| 11 | 2003003 | 0.67 | | | A612E | 0.06 | 0.90 | 0.87 | 0.86 | 0.85 | 0.84 | 0.91 | 0.70 | 1.00 |
| 11 | 2003228 | 0.62 | | | A687E | 0.06 | 0.86 | 0.84 | 0.82 | 0.83 | 0.67 | 0.90 | 0.94 | 1.00 |
| 12 | 73579 | 0.53 | PF3D7_1200900 | Plasmodium exported protein (PHISTc), unknown function | R268. | 0.70 | 0 | 0 | 0 | 0 | 0 | 0 | 0 | 0 |
| 12 | 2118264 | 0.63 | PF3D7_1252100 | rhoptry neck protein 3 | N1004K | 1.00 | 0.24 | 0.16 | 0.28 | 0.48 | 0.94 | 0.69 | 0.95 | 0.60 |
| 13 | 83595 | 0.64 | PF3D7_1301400 | Plasmodium exported protein (hyp12), unknown function | N241K | 0.79 | 0.01 | 0 | 0.01 | 0 | 0 | 0 | 0 | 0 |
| 13 | 92904 | 0.60 | PF3D7_1301600 | erythrocyte binding antigen-140 | H150R | 0.77 | 0.02 | 0 | 0.01 | 0.02 | 0 | 0 | 0 | 0 |
| 13 | 127110 | 0.66 | PF3D7_1302300 | Plasmodium exported protein, unknown function | T11I | 0.96 | 0.17 | 0.10 | 0.18 | 0.08 | 0.05 | 0.01 | 0.39 | 0.93 |
| **13** | **145204** | **0.82** | **PF3D7_1302700** | **ATP-dependent RNA helicase DHR1, putative** | **D11N** | **0.94** | **0.04** | **0.03** | **0.03** | **0** | **0** | **0** | **0** | **0** |
| 13 | 593658 | 0.73 | PF3D7_1313800 | conserved Plasmodium membrane protein, unknown function | H1793L | 0.91 | 0.08 | 0.03 | 0.05 | 0 | 0 | 0 | 0 | 0 |
| 13 | 612670 | 0.53 | PF3D7_1314200 | telomerase reverse transcriptase | Q447K | 0.77 | 0.05 | 0.07 | 0.04 | 0 | 0 | 0 | 0 | 0 |
| 13 | 921466 | 0.51 | PF3D7_1322100 | variant-silencing SET protein | V166I | 0.85 | 0.13 | 0.12 | 0.15 | 0.01 | 0 | 0 | 0 | 0 |
| 13 | 1419303 | 0.60 | PF3D7_1335100 | merozoite surface protein 7 | N280T | 0.89 | 0.18 | 0.06 | 0.12 | 0.22 | 0.15 | 0.12 | 0.19 | 0.56 |
| 13 | 2360342 | 0.59 | PF3D7_1359400 | CUGBP Elav-like family member 1 | A286V | 0.81 | 0.05 | 0.06 | 0.05 | 0.02 | 0 | 0 | 0 | 0 |
| 13 | 2515018 | 0.51 | PF3D7_1362700 | conserved Plasmodium protein, unknown function | N1555S | 0.79 | 0.11 | 0.08 | 0.06 | 0.03 | 0 | 0 | 0 | 0 |
| 13 | 2668749 | 0.50 | PF3D7_1366800 | phosphatidylserine synthase, putative | A298V | 0.98 | 0.34 | 0.22 | 0.32 | 0.03 | 0.01 | 0 | 0 | 0.11 |
| 13 | 2787976 | 0.56 | PF3D7_1370300 | membrane associated histidine-rich protein 1 | A2E | 0.72 | 0 | 0 | 0 | 0 | 0 | 0 | 0 | 0 |

| | | | | | | | | | | | | | |
|---|---|---|---|---|---|---|---|---|---|---|---|---|---|
| 14 | 809757 | 0.70 | PF3D7_1419400 | conserved Plasmodium membrane protein, unknown function | S538F | 0.87 | 0.04 | 0.04 | 0.03 | 0 | 0 | 0 | 0 | 0 |
| 14 | 810165 | 0.69 | | | S402N | 0.87 | 0.05 | 0.05 | 0.03 | 0 | 0 | 0 | 0 | 0 |
| 14 | 823710 | 0.60 | PF3D7_1419800 | glutathione reductase | K72N | 0.83 | 0.07 | 0.06 | 0.04 | 0.01 | 0 | 0 | 0 | 0 |
| 14 | 834826 | 0.62 | PF3D7_1420100 | conserved Plasmodium protein, unknown function | S620N | 0.85 | 0.07 | 0.06 | 0.05 | 0 | 0 | 0 | 0 | 0.02 |
| 14 | 844557 | 0.56 | PF3D7_1420300 | Hsp70-escort protein 1 | V283L | 0.83 | 0.10 | 0.06 | 0.10 | 0.02 | 0 | 0 | 0 | 0 |
| 14 | 2185212 | 0.53 | PF3D7_1453200 | conserved Plasmodium protein, unknown function | S1198Y | 0.85 | 0.25 | 0.04 | 0.10 | 0.63 | 0.04 | 0 | 0 | 0.03 |
| 14 | 2185417 | 0.56 | | | N1230I | 0.87 | 0.22 | 0.06 | 0.11 | 0.06 | 0.28 | 0.64 | 0.09 | 0.02 |
| 14 | 2185421 | 0.51 | | | M1231I | 0.89 | 0.35 | 0.09 | 0.12 | 0.85 | 0.89 | 0.96 | 0.79 | 0.14 |
| 14 | 2186277 | 0.51 | | | K1378N | 0.81 | 0.15 | 0.04 | 0.11 | 0.76 | 0.94 | 0.87 | 0.80 | 0.01 |
| 14 | 2612645 | 0.57 | PF3D7_1464500 | conserved Plasmodium membrane protein, unknown function | E1785G | 0.83 | 0.14 | 0.05 | 0.05 | 0.02 | 0 | 0 | 0 | 0.22 |
| 14 | 2638640 | 0.58 | PF3D7_1465100 | conserved oligomeric Golgi complex subunit 6, putative | I359L | 0.86 | 0.15 | 0.07 | 0.06 | 0.05 | 0.01 | 0.05 | 0 | 0 |
| 14 | 2639315 | 0.52 | | | Y584H | 0.85 | 0.19 | 0.11 | 0.09 | 0.02 | 0.01 | 0.04 | 0 | 0.19 |
| **14** | **2714888** | **0.75** | **PF3D7_1466400** | **AP2 domain transcription factor AP2-EXP** | **L800I** | **0.89** | **0.05** | **0.03** | **0.02** | **0** | **0** | **0** | **0** | **0** |
| 14 | 2948419 | 0.73 | PF3D7_1472200 | histone deacetylase, putative | N920Y | 0.91 | 0.08 | 0.05 | 0.03 | 0.02 | 0.01 | 0.05 | 0 | 0 |
| 14 | 3046529 | 0.51 | **PF3D7_1474400** | **conserved Plasmodium protein, unknown function** | T499A | 0.70 | 0.02 | 0.02 | 0.02 | 0.02 | 0 | 0 | 0 | 0 |
| **14** | **3054932** | **0.88** | | | **I2902N** | **0.97** | **0.05** | **0.03** | **0.02** | **0.01** | **0** | **0.01** | **0** | **0.03** |
| **14** | **3082419** | **0.86** | **PF3D7_1475200** | **conserved protein, unknown function** | **E224K** | **0.96** | **0.05** | **0.02** | **0.03** | **0** | **0** | **0** | **0** | **0.03** |
| 14 | 3088662 | 0.74 | **PF3D7_1475400** | **cysteine repeat modular protein 4** | A5374S | 0.91 | 0.04 | 0.03 | 0.10 | 0 | 0 | 0 | 0 | 0 |
| 14 | 3088877 | 0.74 | | | I5302K | 0.91 | 0.04 | 0.04 | 0.08 | 0 | 0 | 0 | 0 | 0 |
| **14** | **3097632** | **0.75** | | | **Q2384E** | **0.90** | **0.03** | **0.03** | **0.03** | **0** | **0** | **0.01** | **0** | **0** |
| **14** | **3183699** | **0.90** | **PF3D7_1477400** | **Plasmodium exported protein (PHIST), unknown function** | **H87Y** | **0.95** | **0.01** | **0** | **0** | **0** | **0** | **0** | **0** | **0** |
| 14 | 3193593 | 0.59 | PF3D7_1477600 | surface-associated interspersed protein 14.1 (SURFIN 14.1) | K103N | 0.97 | 0.19 | 0.20 | 0.28 | 0.28 | 0.17 | 0.22 | 0.28 | 0.30 |
| 14 | 3193676 | 0.51 | | | R131K | 0.98 | 0.30 | 0.25 | 0.31 | 0.23 | 0.25 | 0.50 | 0.26 | 0.26 |
| 14 | 3193843 | 0.66 | | | L187I | 0.95 | 0.12 | 0.15 | 0.16 | 0.10 | 0.12 | 0.05 | 0.06 | 0 |
| 14 | 3194436 | 0.50 | | | N325D | 0 | 0.66 | 0.64 | 0.70 | 0.56 | 0.72 | 0.81 | 0.75 | 0.97 |

**Supplementary Table 2 – Correlation between AF1 characteristic loci.**

A symmetrical matrix showing the $r^2$ measure of linkage disequilibrium between selected SNPs at six AF1 characteristic loci; each locus is represented by the single SNP with the highest mean $r^2$ with respect to the remaining five loci. The SNP coordinates of each SNP are shown in the form "*chr*:*position*". The remaining columns show: The ID and description of the gene containing the SNP; whether the SNP is nonsynonymous or synonymous, and the amino acid change if any; the mean $F_{ST}$ between AF1 and each of WAF, CAF and EAF populations; and the allele frequency of the AF1 allele in each of the populations, followed by the frequency in other populations present in the Pf7 dataset (SAS=South Asia; WSEA=Western Greater Mekong Subregion; ESEA=Eastern Greater Mekong Subregion; OCE=Oceania; SAM=South America).

| | 02:814192 | 09:1205151 | 10:1413597 | 11:1984241 | 13:2787976 | 14:3183699 | **Variant** | | | | **$F_{ST}$** | **Allele Frequencies** | | | | | | | | |
|---|---|---|---|---|---|---|---|---|---|---|---|---|---|---|---|---|---|---|---|---|
| | | | | | | | **Gene ID** | **Gene Description** | **N/S** | **Amino** | | **AF1** | **WAF** | **CAF** | **EAF** | **SAS** | **WSEA** | **ESEA** | **OCE** | **SAM** |
| 02:814192 | - | 0.47 | 0.51 | 0.53 | 0.51 | 0.43 | PF3D7_0220300 | Plasmodium exported protein, unknown function | N | P92A | 0.56 | 0.72 | 0 | 0 | 0.01 | 0 | 0 | 0 | 0 | 0 |
| 09:1205151 | 0.47 | - | 0.55 | 0.63 | 0.54 | 0.45 | PF3D7_0930300 | merozoite surface protein 1 | N | N1114Y | 0.91 | 0.96 | 0.01 | 0 | 0 | 0 | 0 | 0 | 0 | 0.1 |
| 10:1413597 | 0.51 | 0.55 | - | 0.63 | 0.65 | 0.55 | PF3D7_1035700 | duffy binding-like merozoite surface protein | N | G133D | 0.79 | 0.88 | 0 | 0 | 0 | 0 | 0 | 0 | 0 | 0 |
| 11:1984241 | 0.53 | 0.63 | 0.63 | - | 0.77 | 0.59 | PF3D7_1149300 | serine/threonine protein kinase, FIKK family | S | 559K | 0.73 | 0.85 | 0 | 0 | 0 | 0 | 0 | 0 | 0 | 0 |
| 13:2787976 | 0.51 | 0.54 | 0.65 | 0.77 | - | 0.44 | PF3D7_1370300 | membrane associated histidine-rich protein 1 | N | A2E | 0.56 | 0.72 | 0 | 0 | 0 | 0 | 0 | 0 | 0 | 0 |
| 14:3183699 | 0.43 | 0.45 | 0.55 | 0.59 | 0.44 | - | PF3D7_1477400 | Plasmodium exported protein (PHIST), unknown function | N | H87Y | 0.9 | 0.95 | 0.01 | 0 | 0 | 0 | 0 | 0 | 0 | 0 |

**Supplementary Table 3 – High-IBD genomic regions in the AF1 group**

Each row in this table represents one genome region where ≥50% of AF1 sample pairs are in IBD. From left to right, the columns show: the chromosome number, start position, end position and size of the region; the highest proportion of AF1 sample pairs in IBD in this region, and the position where it occurs; the highest mean $F_{ST}$ (vs the WAF, CAF and EAF populations) in this region, the position where it occurs, the ID and description of the gene, whether the change is synonymous or not, the mutation caused, and the frequency of the non-reference allele in the AF1 population. The regions are sorted in descending order of mean $F_{ST}$. All positions are with respect to the 3D7 v3 reference genome.

| | Region | | | Max IBD pairs | | Maximum $F_{ST}$ Position | | | | | | |
|---|---|---|---|---|---|---|---|---|---|---|---|---|
| Chr | Start | End | Size | % pairs | Pos | $F_{ST}$ | Pos | ID | Description | N/S | Mut | AF1Freq |
| 10 | 1335368 | 1571788 | 236421 | 95.7% | 1442027 | 0.94 | 1456571 | PF3D7_1036900 | conserved Plasmodium protein, unknown function | N | S479I | 0.97 |
| 9 | 1171854 | 1226411 | 54558 | 85.7% | 1202410 | 0.93 | 1205284 | PF3D7_0930300 | merozoite surface protein 1 | N | V1158E | 0.97 |
| 14 | 3183284 | 3207686 | 24403 | 87.5% | 3183284 | 0.90 | 3183699 | PF3D7_1477400 | Plasmodium exported protein (PHIST), unknown function | N | H87Y | 0.95 |
| 9 | 1378245 | 1455635 | 77391 | 95.4% | 1427653 | 0.89 | 1427697 | PF3D7_0936000 | ring-exported protein 2 | N | S77. | 0.98 |
| 14 | 3044149 | 3114489 | 70341 | 85.0% | 3080648 | 0.88 | 3054932 | PF3D7_1474400 | conserved Plasmodium protein, unknown function | N | I2902N | 0.97 |
| 4 | 92597 | 124976 | 32380 | 88.4% | 110070 | 0.85 | 103881 | PF3D7_0401800 | Plasmodium exported protein (PHISTb), unknown function | N | K515R | 0.93 |
| 2 | 299208 | 342734 | 43527 | 60.2% | 306058 | 0.84 | 306406 | PF3D7_0207600 | serine repeat antigen 5 | N | K159E | 0.93 |
| 11 | 1934420 | 2003312 | 68893 | 84.1% | 1985877 | 0.83 | 2001089 | PF3D7_1149600 | DnaJ protein, putative | N | Y8N | 0.96 |
| 13 | 83595 | 167851 | 84257 | 67.2% | 121740 | 0.82 | 145204 | PF3D7_1302700 | ATP-dependent RNA helicase DHR1, putative | N | D11N | 0.94 |
| 1 | 558230 | 569858 | 11629 | 87.2% | 558230 | 0.80 | 563776 | PF3D7_0114700 | PIR protein | N | A300V | 0.93 |
| 2 | 769929 | 814641 | 44713 | 79.6% | 769929 | 0.79 | 784067 | PF3D7_0219700 | gametocyte exported protein 20 | N | Y182H | 0.89 |
| 9 | 777971 | 807125 | 29155 | 73.0% | 781952 | 0.75 | 781952 | PF3D7_0919000 | nucleosome assembly protein | N | I76V | 0.87 |
| 14 | 803248 | 836620 | 33373 | 75.7% | 803248 | 0.70 | 809757 | PF3D7_1419400 | conserved Plasmodium membrane protein, unknown function | N | S538F | 0.87 |
| 8 | 1311766 | 1345474 | 33709 | 80.5% | 1311766 | 0.69 | 1311901 | PF3D7_0830800 | surface-associated interspersed protein 8.2 (SURFIN 8.2) | N | P422R | 0.95 |
| 2 | 109623 | 153841 | 44219 | 56.2% | 109623 | 0.65 | 153549 | PF3D7_0203100 | protein kinase, putative | N | E1145K | 0.80 |
| 3 | 841714 | 866401 | 24688 | 52.6% | 848289 | 0.63 | 865666 | PF3D7_0320700 | signal peptidase complex subunit 2 | N | M78L | 0.89 |
| 14 | 2585522 | 2672686 | 87165 | 67.5% | 2635842 | 0.62 | 2668129 | PF3D7_1465800 | dynein beta chain, putative | S | 1710C | 0.81 |
| 7 | 705498 | 718468 | 12971 | 60.0% | 708260 | 0.59 | 714223 | PF3D7_0716200 | PDCD2 domain-containing protein, putative | S | 585G | 0.77 |
| 6 | 847157 | 888840 | 41684 | 67.4% | 847157 | 0.58 | 851783 | PF3D7_0620400 | merozoite surface protein 10 | N | K391N | 0.89 |
| 5 | 55996 | 128576 | 72581 | 60.8% | 64857 | 0.56 | 93127 | PF3D7_0501800 | CAF-1 p150 homolog | S | 589T | 0.87 |
| 11 | 1856542 | 1866942 | 10401 | 54.3% | 1856542 | 0.53 | 1862558 | PF3D7_1147000 | sporozoite asparagine-rich protein | N | D1600E | 0.70 |
| 11 | 123841 | 138994 | 15154 | 58.6% | 137023 | 0.52 | 123946 | PF3D7_1102600 | gametocyte exported protein 14 | N | L230I | 0.75 |
| 10 | 653023 | 682454 | 29432 | 55.6% | 669207 | 0.51 | 653563 | PF3D7_1016300 | glycophorin binding protein | N | R2Q | 0.71 |

### SUPPLEMENTARY FIGURES

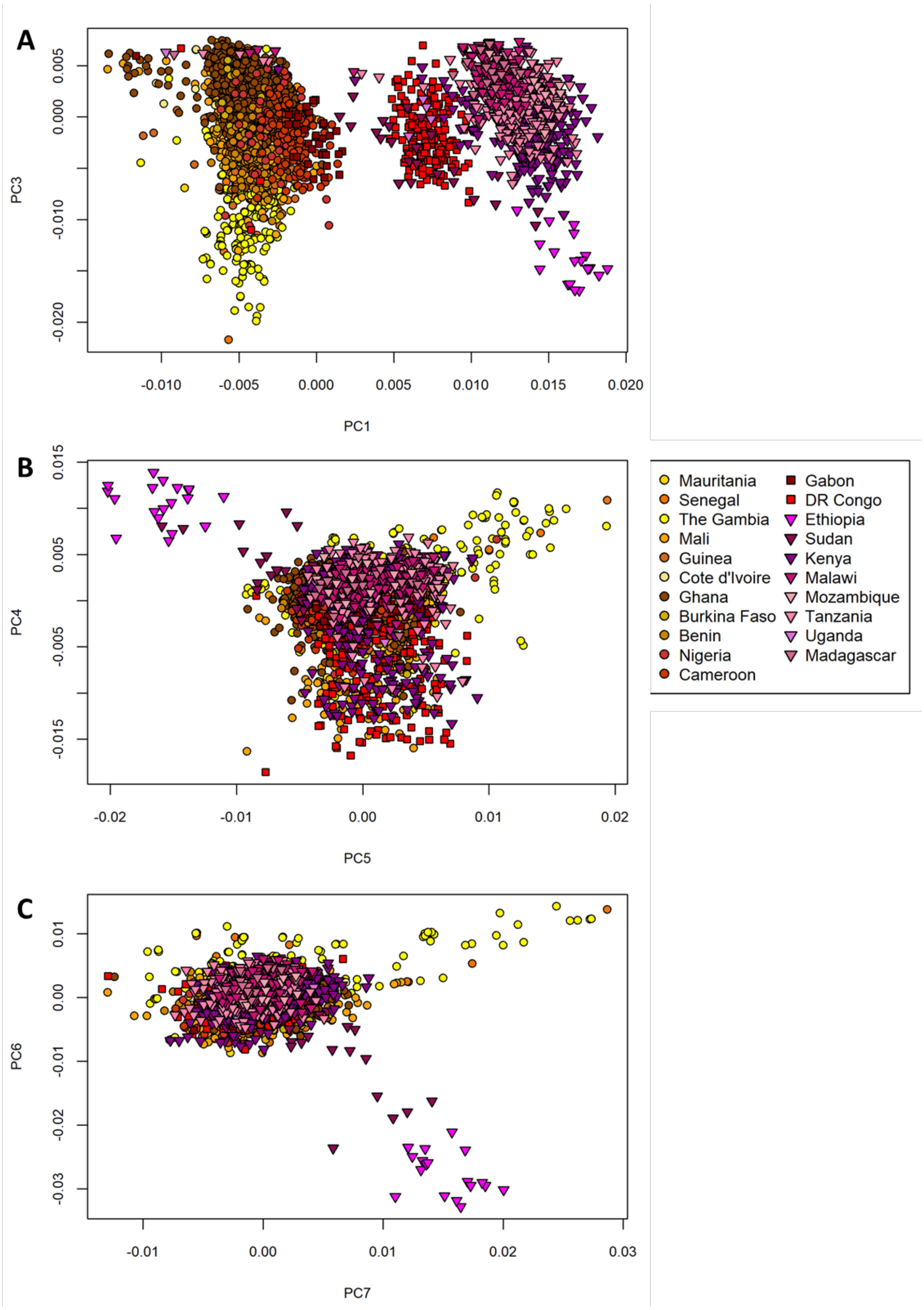

**Supplementary Figure 1 – Plots of higher-order principal components.**

This figure shows higher-order (less significant) principal components in PCoA analysis, from PC3 to PC7: (A) plot of PC1 vs PC3; (B) plot of PC4 vs PC5; and (C) Plot of PC6 vs PC7. All components from PC3 to PC7 appear to be driven by differentiated clusters from Ethiopia (fuchsia-coloured triangle markers) or from the Gambia (yellow circles).

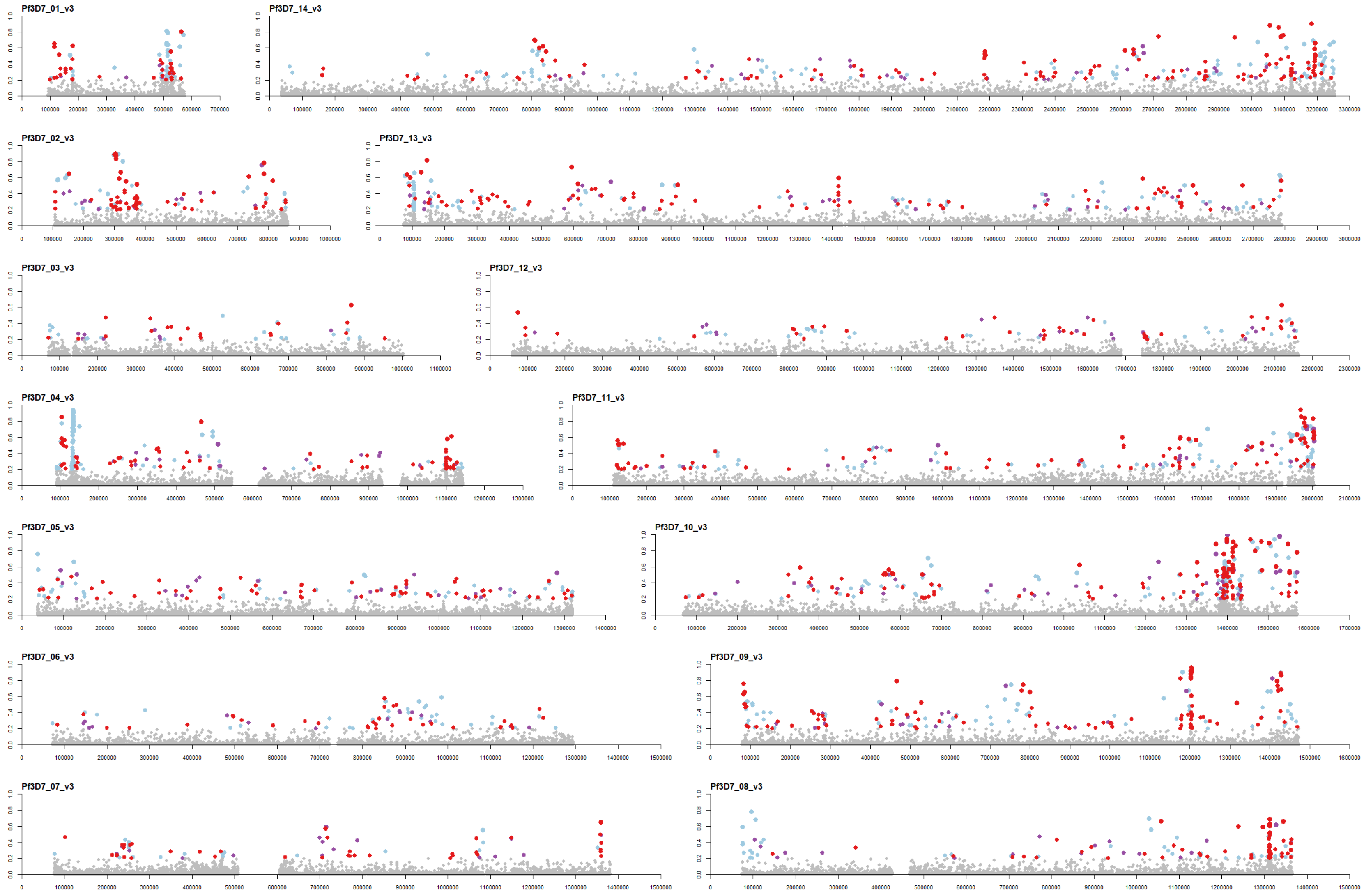

**Supplementary Figure 2 – Genome-wide map of $F_{ST}$ between AF1 and other African populations.**

These plots (one per chromosome, as labelled in the upper left-hand corner of each plot) show the mean $F_{ST}$ between AF1 and the three African macro-regions (WAF, CAF and EAF) at 743,583 SNPs. At each position, we plotted the $F_{ST}$ value (between 0 and 1). Positions with $F_{ST}$≥0.2 are shown by markers coloured according to the type of SNP: light blue for non-coding, purple for synonymous coding, and red for non-synonymous coding variants; SNPs with $F_{ST}$<0.2 are shown by gray markers.

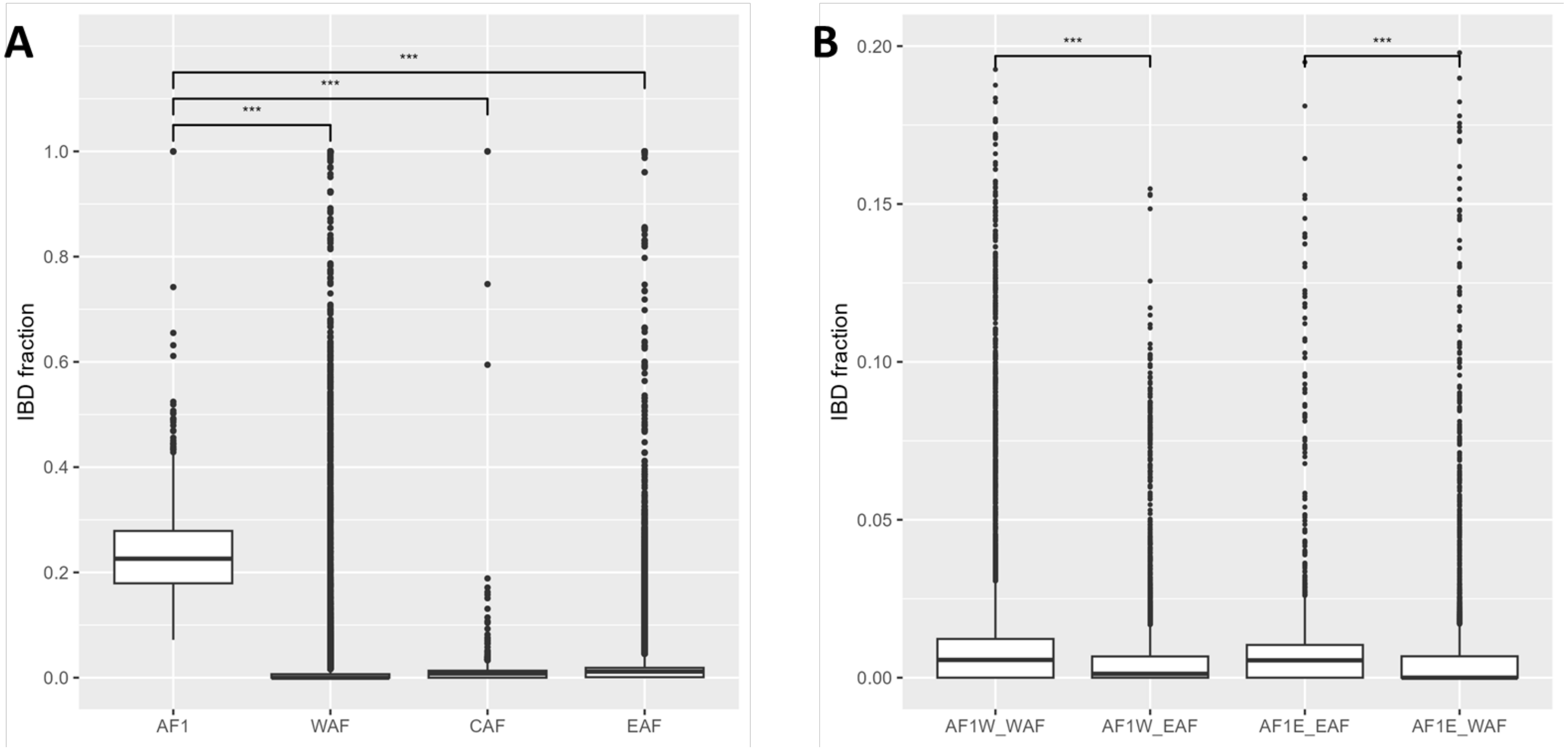

**Supplementary Figure 3 – Pairwise IBD fraction levels within and between African populations.**

(A) Boxplot showing the distribution of IBD genome fractions between all pairs of parasites in each of four populations: AF1, West Africa (WAF), Central Africa (CAF) and East Africa (EAF). Levels within AF1 are significantly higher than within the rest of the populations ($p<0.001$). (B) Boxplot showing the distribution of IBD genome fractions between populations. The first two columns show IBD fractions for all pairings of West African AF1 members (AF1W) with West African (column 1) and East African (column 2) non-AF1 parasites (WAF and EAF respectively). The remaining two columns show the IBD fractions for all pairings of East African AF1 members (AF1E) with East African (column 3) and West African (column 4) non-AF1 parasites. Although IBD levels between populations are low, they are significantly higher between AF1 members and non-AF1 parasites from the same regions than those from a different region.

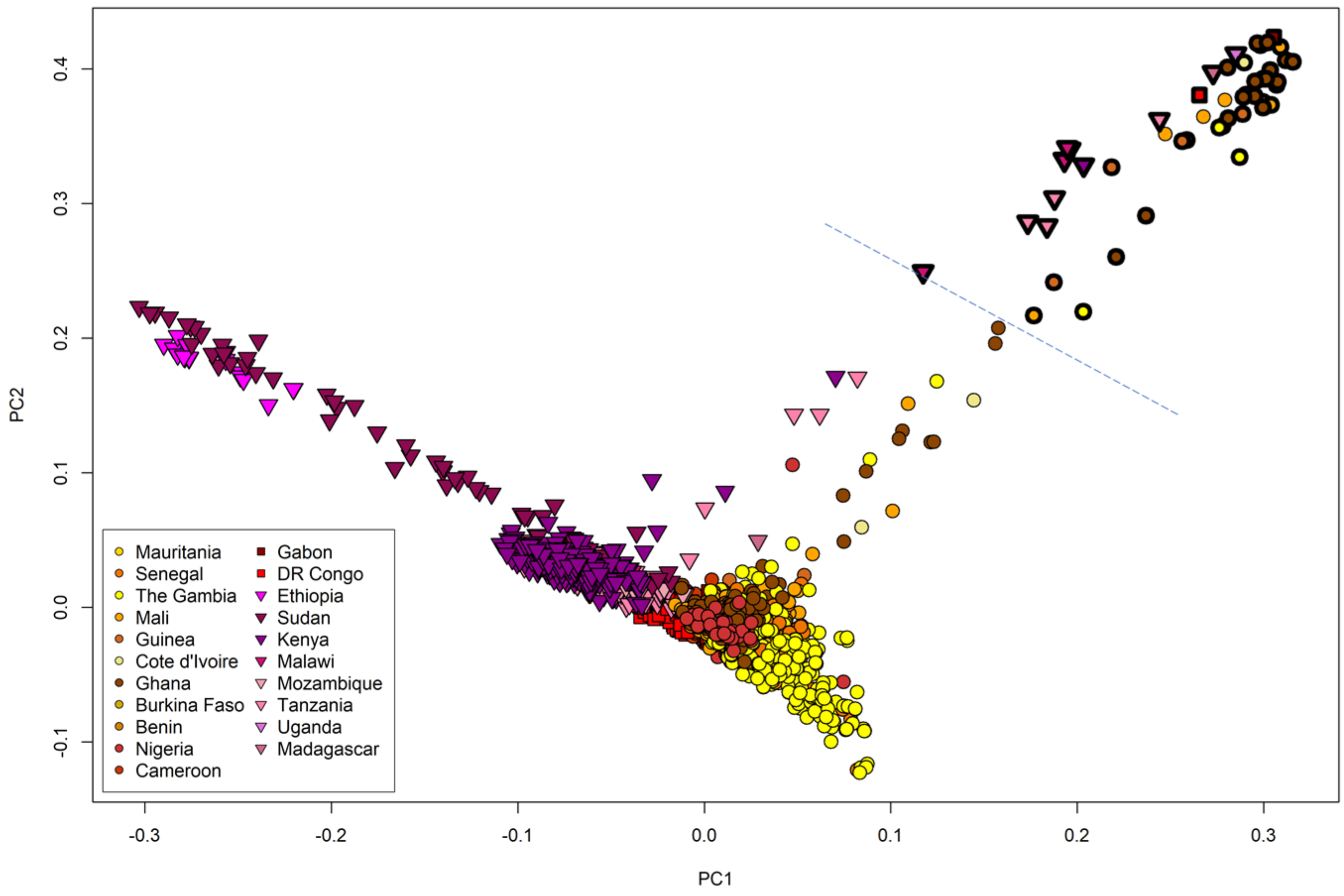

**Supplementary Figure 4 – PCoA plot based on an IBD distance measure.**
This figure shows the first two components of a PCoA using an IBD-based distance measure ($d$=1-$frac$, where $frac$ is the fraction of the genome in pairwise IBD). Samples are coloured by country of origin, and AF1 parasites are shown with a thicker border. A blue dotted line shows a cut-off between AF1 and non-AF1 parasites. Three individuals from Mali (shown as orange circles) have joined the AF1 group; they were found to have substantial genotype missingness, which may have affected their classification.

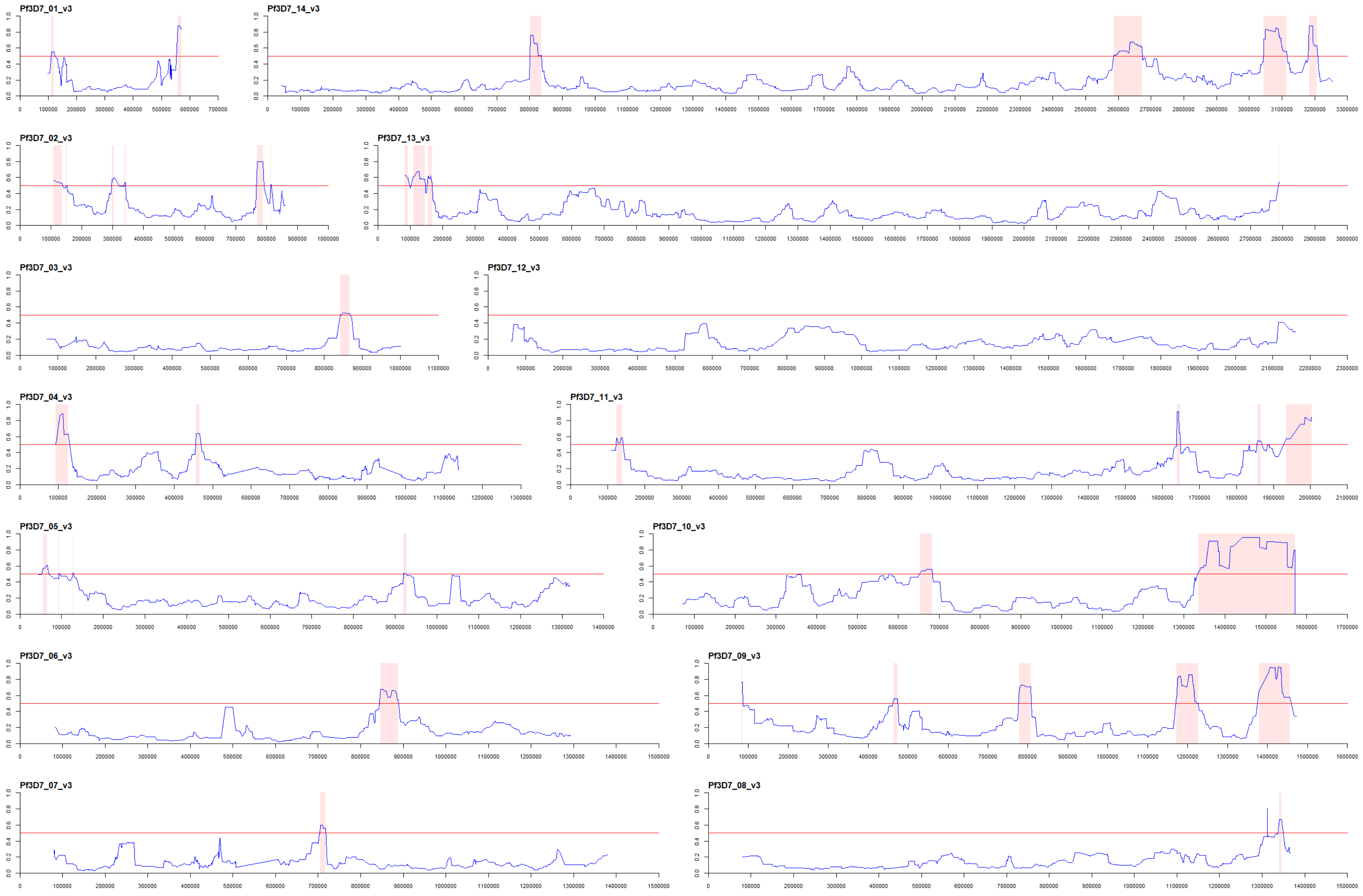

**Supplementary Figure 5 – Genome-wide map of pairwise IBD within the AF1 population.**
The 14 plots (one per chromosome, as labelled in the upper left-hand corner of each plot) show the proportion of AF1 sample pairs that are predicted to be identical by descent at 43,469 SNPs (blue line). Region with ≥50% IBD sample pairs (red threshold line) are highlighted by a pink background.

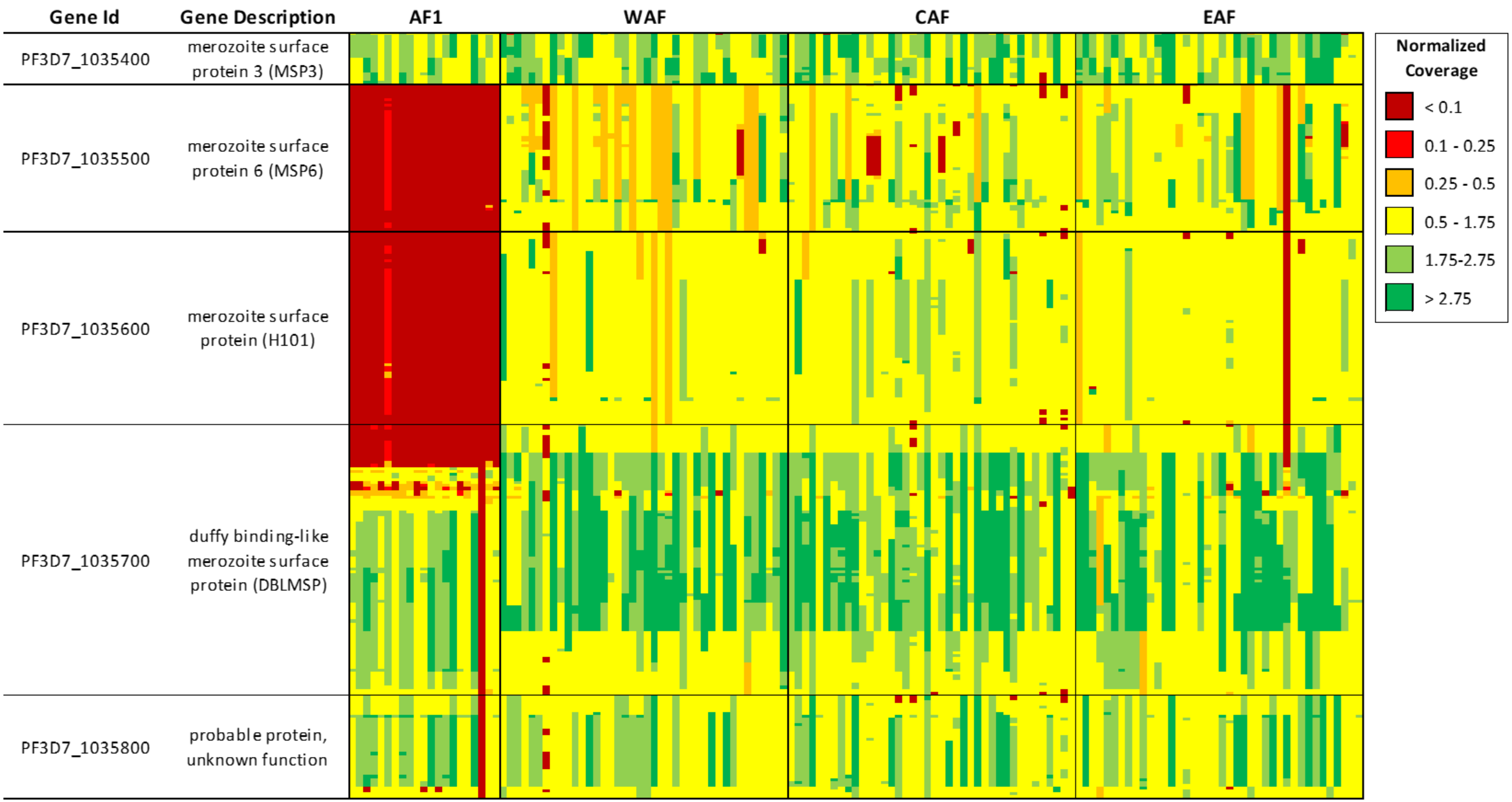

**Supplementary Figure 6 – Coverage analysis of the Chromosome 10 locus.**
This matrix shows sequencing reads coverage at SNPs in the region of the AF1 characteristic locus on chromosome 10. Each row represents a SNP, with the first two columns showing the ID and description of the containing gene; each column represents a sample. Samples that have undergone selective whole-genome amplification (sWGA) were excluded since sWGA affects coverage patterns. All non-sWGA AF1 samples (n=21) are shown, followed by 40 randomly selected samples for each of the African macro-regions (WAF, CAF and EAF) as labelled. Normalized coverage was computed by dividing the number of reads by the genome-wide median coverage in the sample. AF1 samples have extremely low coverage over genes MSP6 and H101 (dark red coverage region), suggesting a large deletion.

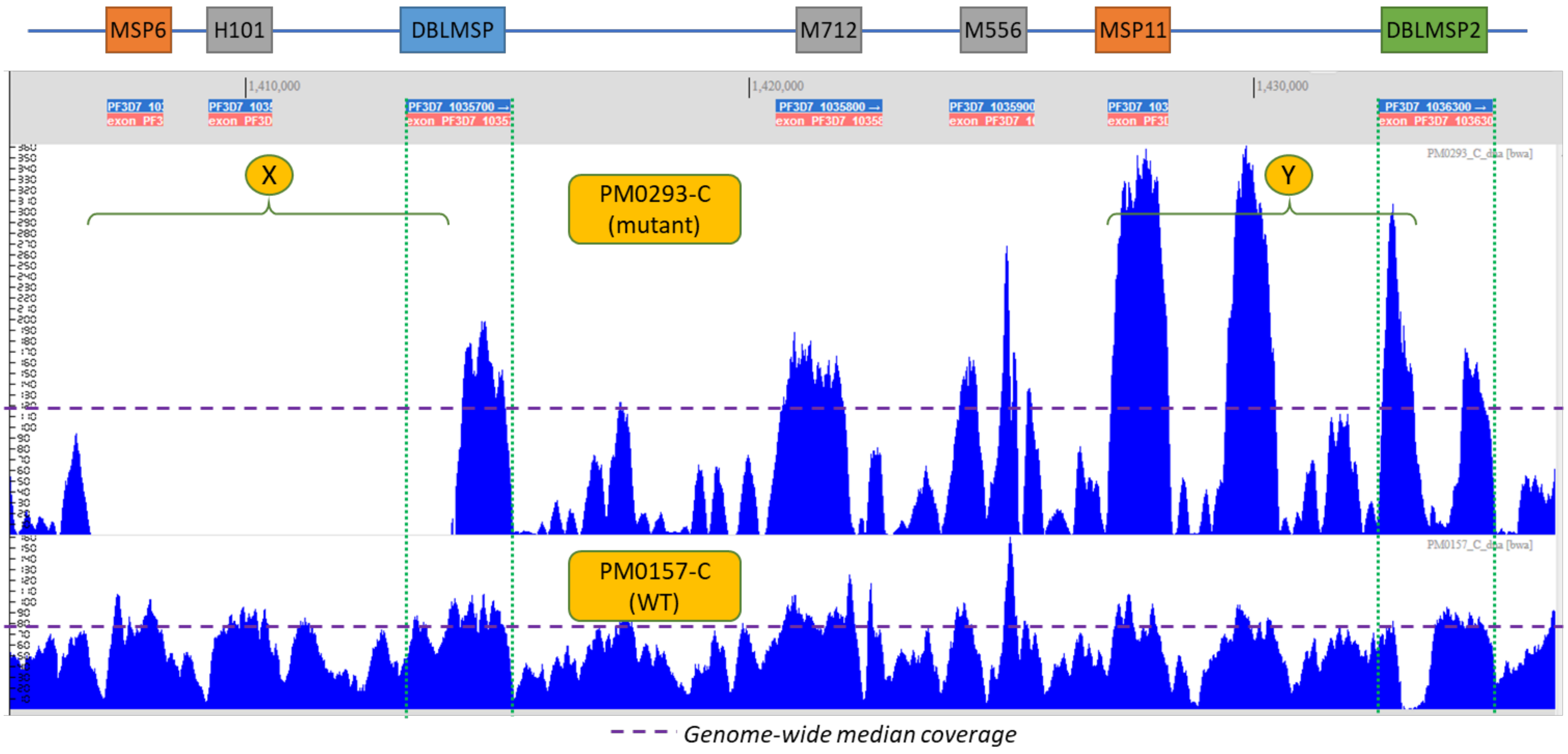

**Supplementary Figure 7 – Coverage of the Chromosome 10 locus in an AF1 sample.**
The above figure shows two pileup plots for the largest AF1 characteristic locus on chromosome 10 (visualized using the LookSeq genome browser[2]). The height of the pileup indicates the read coverage (as shown on the y-axis). The upper plot shows the pileup for reads from the PM0293-C AF1 member from Mali, while the lower plot shows the pileup for PM0157-C, a non-AF1 parasite from Mali. The genome coordinates (relative to the Pf3D7 reference genome) and the extent of the genes covered by this plot are shown above the pileup plots, topped by coloured boxes showing the genes' names. Purple dashed lines show the genome-wide median read coverage for the two samples. Two regions are demarcated: region "X" shows no coverage over genes MSP6 and H101, and over the 5' end of DBLMSP, suggesting a large deletion; and region "Y" shows high coverage over MSP11 and the 5' end of DBLMSP2.

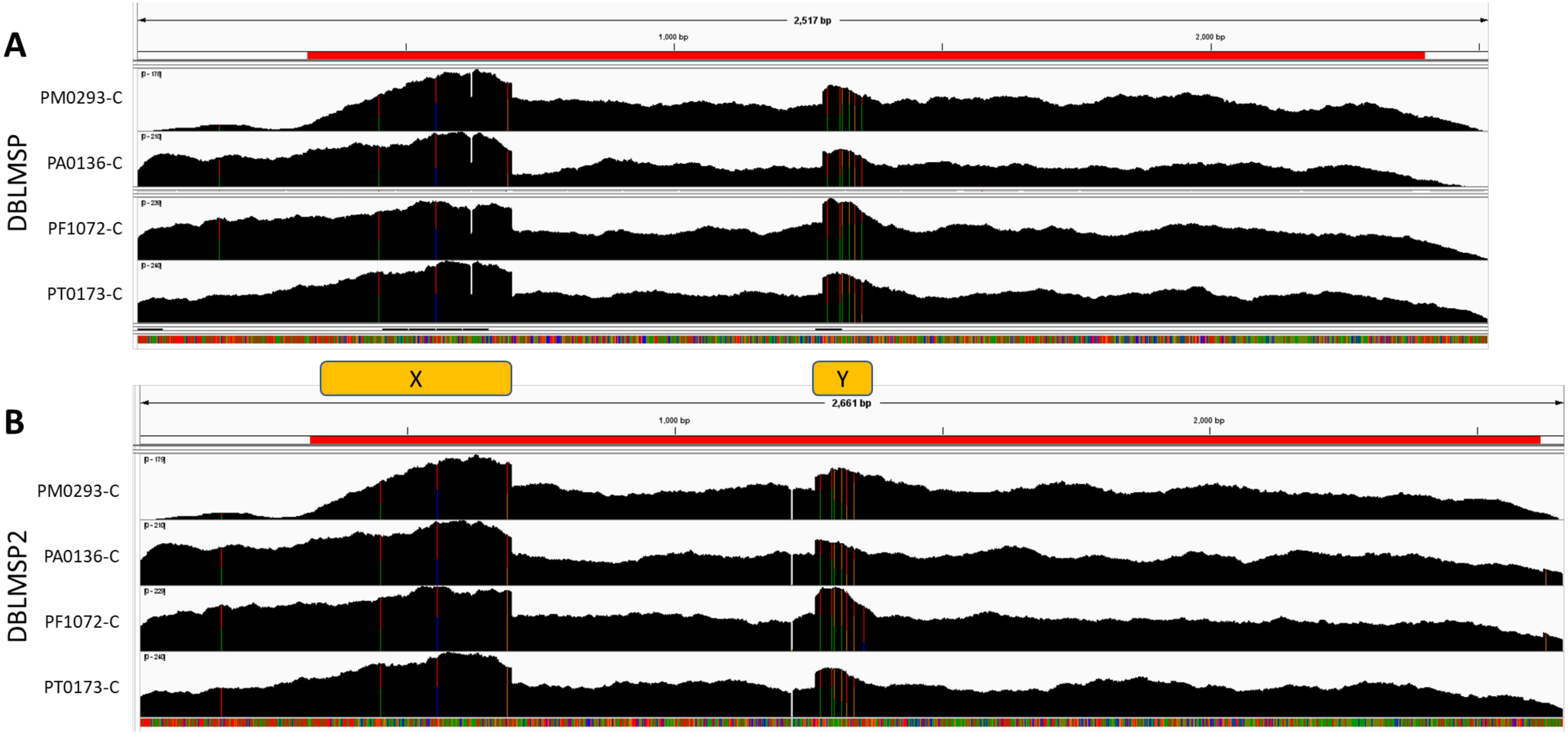

**Supplementary Figure 8 – Coverage profiles of AF1 sequencing read alignments on predicted *de novo* assembly reads.**
The two panels show plots of coverage in alignments of sequencing reads for four AF1 samples (PM0293-C, PA0136-C, PF1072-C and PT173-C) using as references the *de novo* assembled sequences of DBLMSP (panel A) and DBLMSP2 (panel B) from AF1 sample PM0293-C. The red stripe in the upper section of each panel indicates the coding sequence of the gene used as reference. The alignments were conducted separately to avoid alignment competition (i.e. the same reads may have been mapped in both alignments). For both genes, all four samples show fairly even coverage, without sizeable coverage gaps, over most of the coding sequence (in contrast with no coverage in the 5' regions of DBLMSP when aligned against Pf3D7, see Supplementary Figure 7), consistent with a correct assembly of the AF1 sequences. At the 5' end of each alignment (denoted by "X"), there is an approximate doubling in coverage, consistent with the two genes having near-identical sequences in that region, such that reads sequenced from both genes map to the region. A similar effect is observed in the region containing the breakpoint sequence ("Y"). The visualizations were created using the from the Integrative Genomics Viewer (IGV)[1].

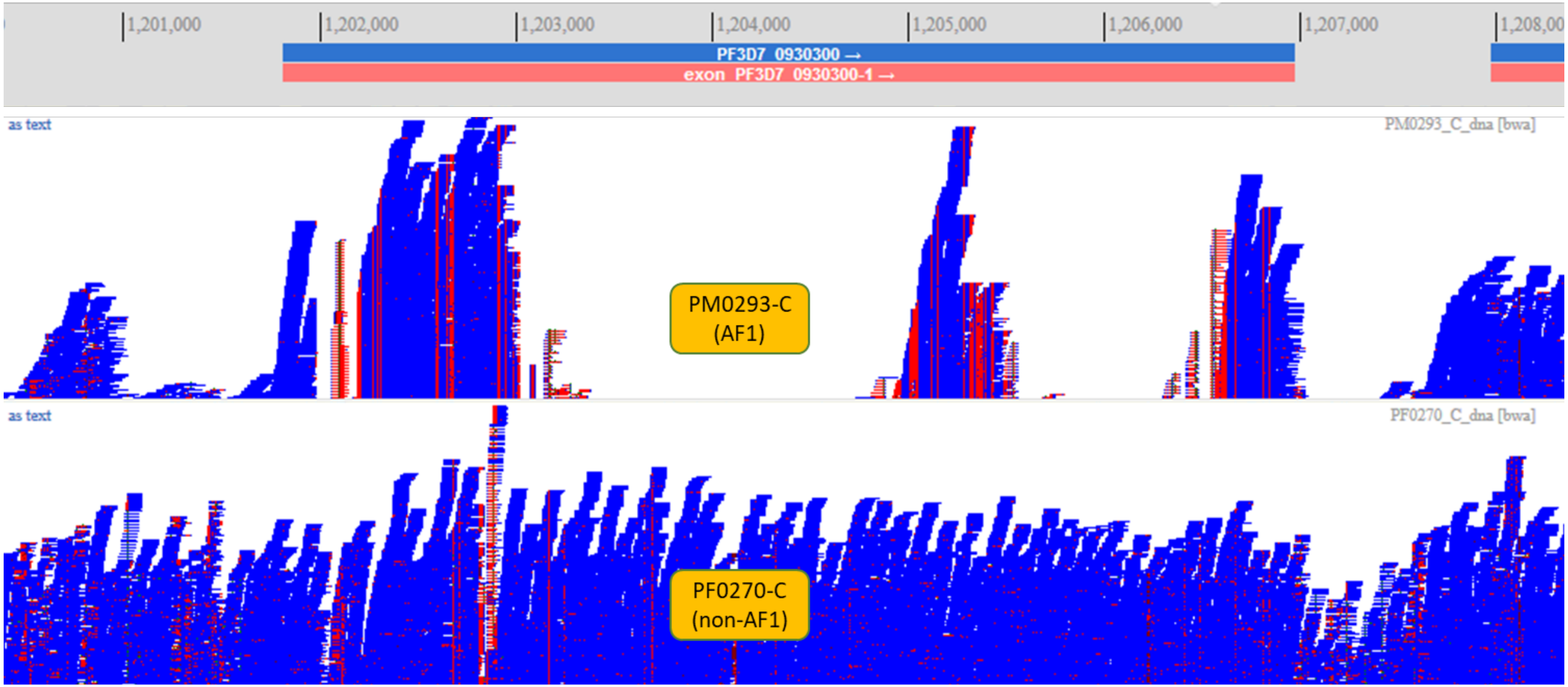

**Supplementary Figure 9 – Alignment of AF1 sequencing reads in the MSP1 gene.**
The above figure shows two pileup plots (visualized using the LookSeq genome browser[2]) in which the height of the pileup indicates the read coverage. The upper plot shows the pileup for reads from the PM0293-C AF1 member from Mali, while the lower plot shows the pileup for PF0270-C, a non-AF1 parasite from Ghana. The genome coordinates (relative to the Pf3D7 reference genome) and the extent of the MSP1 gene (Pf3D7_0930300) are shown above the pileup plots. The large gaps in the AF1 pileup denote regions of the gene (blocks) where the PM0290-C sequence is highly differentiated with respect to the Pf3D7 reference, to the extent that sequencing reads cannot be mapped. This does not occur in the PF0270-C genome, whose sequence is similar to that of Pf3D7.

#### REFERENCES FOR SUPPLEMENTARY MATERIALS

1. Robinson JT, Thorvaldsdottir H, Winckler W, et al. Integrative genomics viewer. *Nat Biotechnol* 2011; **29**(1): 24-6.
2. Manske HM, Kwiatkowski DP. LookSeq: a browser-based viewer for deep sequencing data. *Genome research* 2009; **19**(11): 2125-32.
